# Malaria resistance-related biological adaptation and complex evolutionary footprints of Tai-Kadai people inferred from 796 genomes

**DOI:** 10.1101/2023.07.04.547671

**Authors:** Shuhan Duan, Mengge Wang, Zhiyong Wang, Yan Liu, Xiucheng Jiang, Haoran Su, Yan Cai, Qiuxia Sun, Yuntao Sun, Xiangping Li, Jing Chen, Yijiu Zhang, Jiangwei Yan, Shengjie Nie, Liping Hu, Renkuan Tang, Libing Yun, Chuan-Chao Wang, Chao Liu, Junbao Yang, Guanglin He

**Affiliations:** Institute of Basic Medicine and Forensic Medicine, North Sichuan Medical College and Center for Genetics and Prenatal Diagnosis, Affiliated Hospital of North Sichuan Medical College, Nanchong, Sichuan, 637007, China; Institute of Rare Diseases, West China Hospital of Sichuan University, Sichuan University, Chengdu, 610000, China; Center for Genetics and Prenatal Diagnosis, Affiliated Hospital of North Sichuan Medical College, Nanchong, Sichuan, 637007, China; Department of Forensic Medicine, College of Basic Medicine, Chongqing Medical University, Chongqing, 400331, China; School of Forensic Medicine, Shanxi Medical University, Jinzhong, 030001, China; School of Forensic Medicine, Kunming Medical University, Kunming, 650500, China; West China School of Basic Science & Forensic Medicine, Sichuan University, Chengdu, 610041, China; Faculty of Forensic Medicine, Zhongshan School of Medicine, Sun Yat-sen University, Guangzhou, 510275, China; Guangzhou Forensic Science Institute, Guangzhou, 510055, China; Anti-Drug Technology Center of Guangdong Province, Guangzhou, 510230, China; State Key Laboratory of Cellular Stress Biology, National Institute for Data Science in Health and Medicine, School of Life Sciences, Xiamen University, Xiamen 361005, Fujian, China; Center for Archaeological Science, Sichuan University, Chengdu, 610000, China

**Keywords:** Biological adaptation, genetic admixture, population history, Tai-Kadai people, Malaria resistance

## Abstract

Pathogen-host adaptative interaction and complex population demographical processes, including admixture, drift and Darwen selection, have considerably shaped the Neolithic-to-Modern Western Eurasian population structure and genetic susceptibility to modern human diseases. However, the genetic footprints of evolutionary events in East Asia keep unknown as the underrepresentation of genomic diversity and the design of large-scale population studies. We reported one aggregated database of genome-wide-SNP variations from 796 Tai-Kadai (TK) genomes, including Bouyei first reported here, to explore the genetic history, population structure and biological adaptative features of TK-speaking people from Southern China and Southeast Asia. We found geography-related population substructure among TK-speaking people using the state-of-the-art population genetic structure reconstruction techniques based on the allele frequency spectrum and haplotype-resolved phased fragments. We found that the Northern TK-speaking people from Guizhou harboured one TK-dominant ancestry maximised in Bouyei people, and the Southern one from Thailand obtained more influences from Southeast Asians and indigenous people. We reconstructed the fitted admixture models and demographic graphs, which showed that TK-speaking people received gene flow from ancient rice farmer-related lineages related to the Hmong-Mien and Austroasiatic people and Northern millet farmers associated with the Sino-Tibetan people. Biological adaptation focused on our identified unique TK lineages related to Bouyei showed many adaptive signatures conferring Malaria resistance and low-rate lipid metabolism. Further gene enrichment, the allele frequency distribution of derived alleles, and their correlation with the incidence of Malaria further confirmed that *CR1* played an essential role in the resistance of Malaria in the ancient “Baiyue” tribes.

## INTRODUCTION

East Asia is one of the densest population residence areas in the world and is characterised by abundant cultural, linguistic, and genetic diversity. Ancient East Asian genomes revealed a high degree of genetic differentiation and large-scale population admixture between ancient Northern and Southern East Asians since the early Neolithic period(1). It was common to observe genetic stability or continuity in the Tibetan Plateau, Amur River Basin, Fujian, and Taiwan Island, which differed from the massive migrations and complex admixture scenarios observed in Europe and Southeast Asia(2–5). Agriculture-derived population expansion and migrations shaped the patterns of genetic and linguistic diversity in the core regions of China and Southeast Asia (SEA), supporting the farming-language-people dispersal hypothesis(5, 6). People spread around the Amur River (AR) in Northern China were mainly accompanied by the spread of the Altaic languages(3). Millet farmers dispersed from the Yellow River Basin (YRB) in Central China and this population expansion was associated with the spread of Sino-Tibetan (TB) languages across eastern Eurasia(7). Rice farmers migrated from the Yangtze River Basin, accompanied by the spread of Hmong-Mien (HM), Tai-Kadai (TK), Austroasiatic (AA), and Austronesian (AN) languages in South China(3).

Southern China is the birthplace of rice-cultivating agriculture and is a pivotal crossroads for rice farmers migrating southward to SEA(8). Meanwhile, abundant linguistic, cultural and ethnic diversity contributed to the mysterious verve of the evolutionary history of Southern China(9). TK-speaking populations were the indigenous people of Southern China and widely distributed in Southern China, mainland Southeast Asian (MSEA), and South Asia, ranging from Hainan Island in the East to Northeast India in the West, from Southern Sichuan in the North to Southern Thailand in the South. According to archaeological and historical documents, the ancient “Baiyue” living in Southeast China was considered the ancestors of the present TK-speaking populations(10). During the Han Dynasty, under the pressure of war and famine, numerous “Baiyue” people expanded southward to Southwest China and SEA for long periods. Ancient DNA evidence has illuminated that the Bronze Age migration of farming people had brought TK ancestry and culture to SEA(5, 6). Innumerable subsequent isolation and genetic admixture events further shaped the specific patterns of the genetic structure of present-day TK-speaking populations(11). However, due to the geographical proximity of the TK-speaking people to other Southern Chinese groups (HM, AN and AA), the genetic origins of TK-speaking people, phylogenetic relationships between TK-speaking populations and geographical neighbours and genetic signatures of pathogen-host interaction remained to be fully characterised.

Previous work provided new insights into the population history of Proto-TK-speaking people and their interaction with modern and ancient Southeast Chinese people(12, 13). Elsewise, genetic evidence provided essential clues for the formation of some geographically restricted TK-speaking people in China based on Y-chromosome, mitochondrial and forensic-related low-resolution genetic variations(12, 13). Cultural documents suggested that Hanging Coffin relics in Southern China and SEA shared a lot of common cultural elements with the ancient “Baiyue” tribe relics, further providing cultural evidence for the South China Origin of TK-speaking people. Zhang et al. reported that the ancient Southern Chinese populations associated with Hanging Coffin originated from the coastal region of Southern China (likely in the Mount Wuyi region of China) about 3,600 YBP based on the maternal genetic evidence(14). Wang et al. identified two historic populations in Guangxi strongly associated with modern linguistically-different people: 1500-year-old BaBanQinCen (BBQC) related to TK speakers and 500-year-old Gaohuahua connected to HM speakers(15). Contemporary genetic evidence also produced new insights into the population formation of TK-speaking people. He et al. found an excellent representative source for TK-speaking people on Hainan Island(13). Chen et al. investigated the admixture history of Hainan Li people based on the whole-genome sequence (WGS) data and illuminated that TK-speaking populations from South China and North Vietnam showed close genetic affinity and had a common genetic origin of geographically different “Baiyue” lineages(16). Besides, this work also estimated the formation time of the Li-specific lineage O-M95 and refined the possible origin time of the “Baiyue” lineage about ∼11,000 years ago(16). Nevertheless, the lack of systematic research about the genetic substructure of TK in inland China still hindered our understanding of the whole landscape of TK-speaking people. Preceding genetic analyses were mainly focused on single inland TK-speaking populations or geographically restricted groups based on low-density forensic genetic markers (such as traditional Y and mtDNA genetic markers) or overlapping low-density 50K SNPs(17–22). Therefore, intensive and depth genetic studies focused on the evolutionary features of TK-speaking people are needed. More efforts should be paid to focus on multi-regional integration, systematic scale description, complex population modelling, and biological adaptative, especially the mountain population of Bouyei. The Bouyei are an ethnic group mostly living in Guizhou Province(23). Guizhou has complex landforms and numerous mountain ranges. It is an essential part of the Yungui Plateau and is geographically close to Yunnan and Guangxi provinces, which possess substantial sociocultural, genetic, and linguistic diversities(24). There are more than 20 officially recognised or unrecognised ethnic groups widely distributed in Guizhou, and the complexity of geographical environments and ecological diversities further provide favourable conditions for forming the unique genetic structure of these ethnic groups. Bouyei is one of the 18 officially recognised ethnic minorities in Guizhou Province, which is also distributed in Yunnan, Sichuan, and other provinces. Among these, Guizhou has the largest Bouyei population, accounting for about 97% of the Bouyei population in China. The national language of Bouyei belongs to the TK (also known as Krai-Dai) language family, and their ancient ancestors inhabited Guizhou Province since the Stone Age and grew rice and other crops for a living. Ren et al. made a preliminary exploration focused on the genetic structure of Bouyei populations in Guizhou based on the short tandem repeats (STRs) on X chromosomes(25). He et al. also explored Bouyei’s genetic diversity and forensic characteristics based on insertion/deletion markers(12). Previous genetic studies on the exploration of forensic characteristics and population admixture of Bouyei groups were primarily based on low-density genetic markers, while Bouyei’s fine-scale genetic structure and detailed genetic history remained unclear. Thus, we made a comprehensive population genetic analysis to describe their ancestral composition and reveal their genetic origin. To systematically analyse the genetic structure, evolutional history and environmental adaption of geographically different TK-speaking populations, we newly collected Bouyei samples in Guizhou Bannong (BBN) and merged our data with publicly available array-based genotyping data of modern and ancient Eurasian populations. In this study, we would like to shed light on the following four questions:

1. What are the overall patterns of the genetic diversity of TK-speaking people, and how about the impact of the geographical and cultural factors on it;
2. How many ancestral sources contributed to the gene pool of modern TK-speaking people, and what role of the Bouyei-related ancestry play in it?
3. How about the genetic homogeneity and heterozygosity among ethnically different TK-speaking people from South China and SEA and geographically different Bouyei people inferred from the shared fine-scale haplotypes and allele frequency spectrum?
4. What are the influences of the selection pressures on the genetic architecture of TK-speaking people, and how about the genetic legacy of their interaction with ancient Malaria-related infectious disease? We provide new insights into the genetic admixture history of TK-speaking people (especially the newly-reported Bouyei-related ancestry) and inferred signatures of natural selection in mountainous circumstances based on 796 TK genomes.

## Results

### General patterns of population structure of TK-speaking people in the context of worldwide populations

To dissect the ancestral components and genetic similarity of TK-speaking people, we conducted a model-based ADMIXTURE analysis among 207 modern worldwide populations from our previous published data and reference data from HGDP and Oceania genomic resources(26, 27). When K=11, we observed five East Asian-related ancestries [Yakut-related (blue), Yao-related (orange), Hui-related (light blue), Tibetan-related (light orange) and Bouyei-related (Red)] and six non-East Asian ancestries [Papuan-related (dark green), Solomen-related (dark blue), Sardinian-related (light purple), Karitiana/Surui-related (light green), Mbuti-related (brown), Kalash-related (light brown)] (**Figure 1A**). The red ancestry component widely existed in our studied Bouyei and their neighbours, which was first reported as inland TK ancestry in our work. Bouyei had the highest proportion of this newly-identified ancestry (79.5%±0.0885), followed by Yao (9%) and Hui (7.5%) from South China, and Tibetan had less inland TK-related ancestry but could not be ignored, which suggested that inland TK-related ancestry played an essential role in the formation of ancient Bouyei people and modern southern Chinese people. Other TK-speaking people have similar admixture patterns with different proportions (**Figure S1**). Notably, the ancestral proportions of Northern East Asians (Tibetan-related component) in some TK-speaking populations were remarkable, especially in Gelao, which was markedly different from the ancestral composition patterns observed in our studied groups (**Figure S1**).

**Figure 1.**
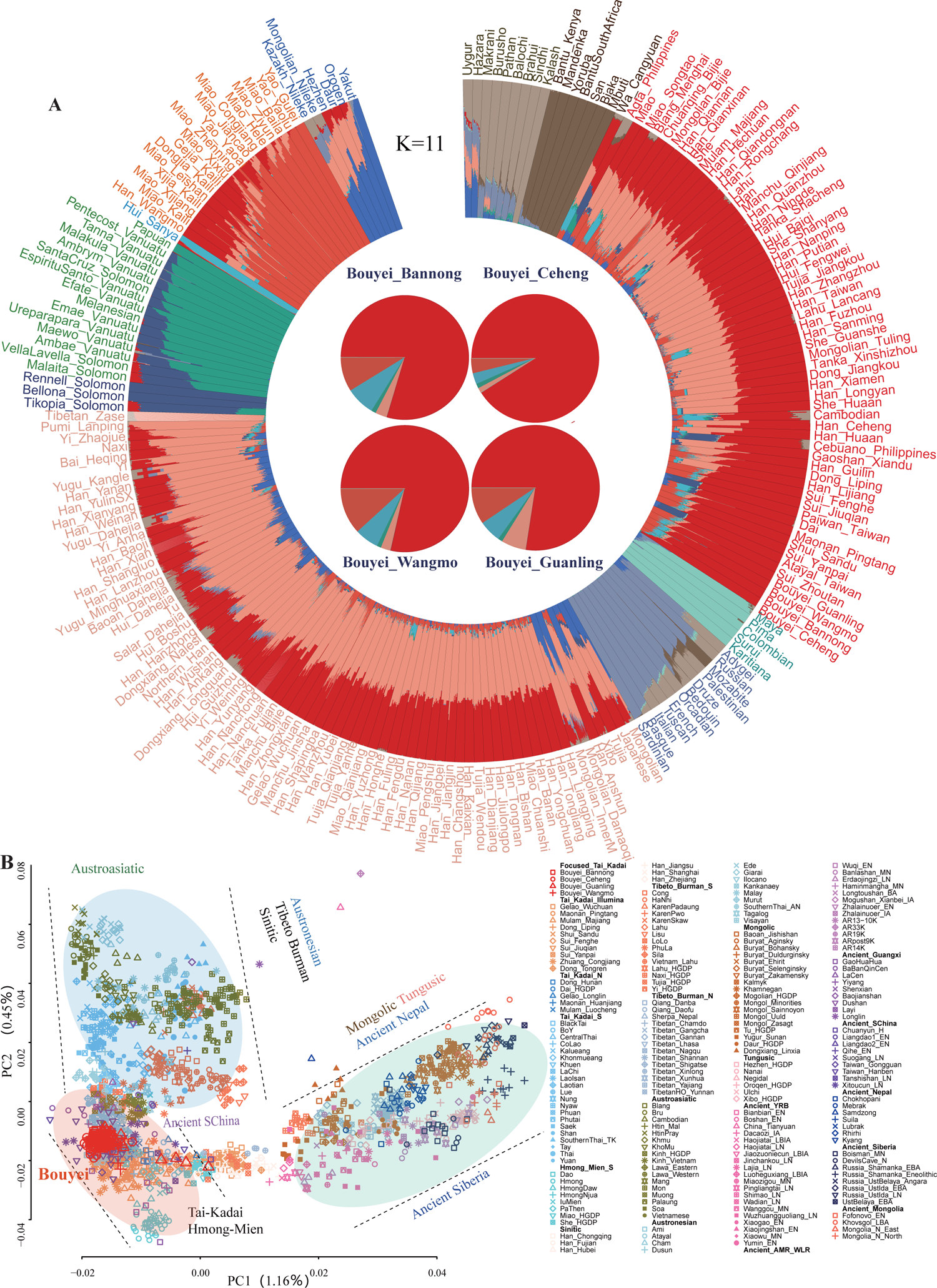
General genetic affinity and population structure from different populations. (**A**) Results of model-based ADMIXTURE clustering analysis. Clustering patterns were visualised with the 207 populations at K=11 based on a high-density dataset. All of these ancestral components were presented by different colours. (**B**) Patterns of genetic relationship based on low-density merged HO dataset: 207 East Asian populations-based PCA was conducted based on the genetic variations of modern and ancient people were projected onto it. (**C**) Populations were colour-coded based on their language family categories, and different shapes represented different populations.

To explore the genetic affinity between Chinese TK-speaking people and southern TK-speaking people from Vietnam, Thailand and Laos (**Figure 1B**), we merged our genome-wide data with publicly available modern and ancient people genotyped via Affymetrix Human Origin array and formed the second dataset, which was referred to as the low-density merged HO **(Table S1, Figure 1C**). We conducted Principal Components Analysis (PCA) with ancient people projected onto modern people’s genetic backgrounds and identified three clines related to Northern Altaic cline, Central HM/TK cline and Southern AA/AN/TK-related cline. The first component extracted 1.16% variance and distinguished HM and TK speakers from Tungusic and Mongolic populations in North China and Ancient Siberians. The second component (PC2: 0.45%) separated HM/TK speakers from AN/AA/TK-speaking people. Ancient people from Guangxi, including historic Gaohuahua and BBQC, were clustered closely with Chinese HM and TK-speaking people, respectively.

Furthermore, we explored the admixture history of TK-speaking populations in the context of modern and ancient Southern East Asia. We observed that TK-speaking populations mainly gathered with HM-speaking populations and were far away from AA/AN/TB-speaking populations, they also showed a close genetic similarity with ancient people from Guangxi, Fujian and surrounding regions, which implied that the ancestors of TK-speaking populations were possibly relevant to the descendants of Southern Chinese (**Figure 2A**). Moreover, we focused on the demographical events that occurred within 39 TK-speaking populations from MSEA and Southern China to explore the fine-scale genetic affinities and their influence on linguistically different populations (**Table S2, Figure S2**). PCAs based on linkage disequilibrium (LD)-independent SNPs showed that the dispersion of TK-speaking populations was related to their geographical locations (**Figure 2B**). The ADMIXTURE results of 117 Southern Chinese populations with K=6 showed that TK-related component gradually decreased from North to South, and TK-speaking populations from MSEA derived more ancestral components from AA-related ancestry compared with TK-speaking populations from China (**Figure 2C**).

**Figure 2.**
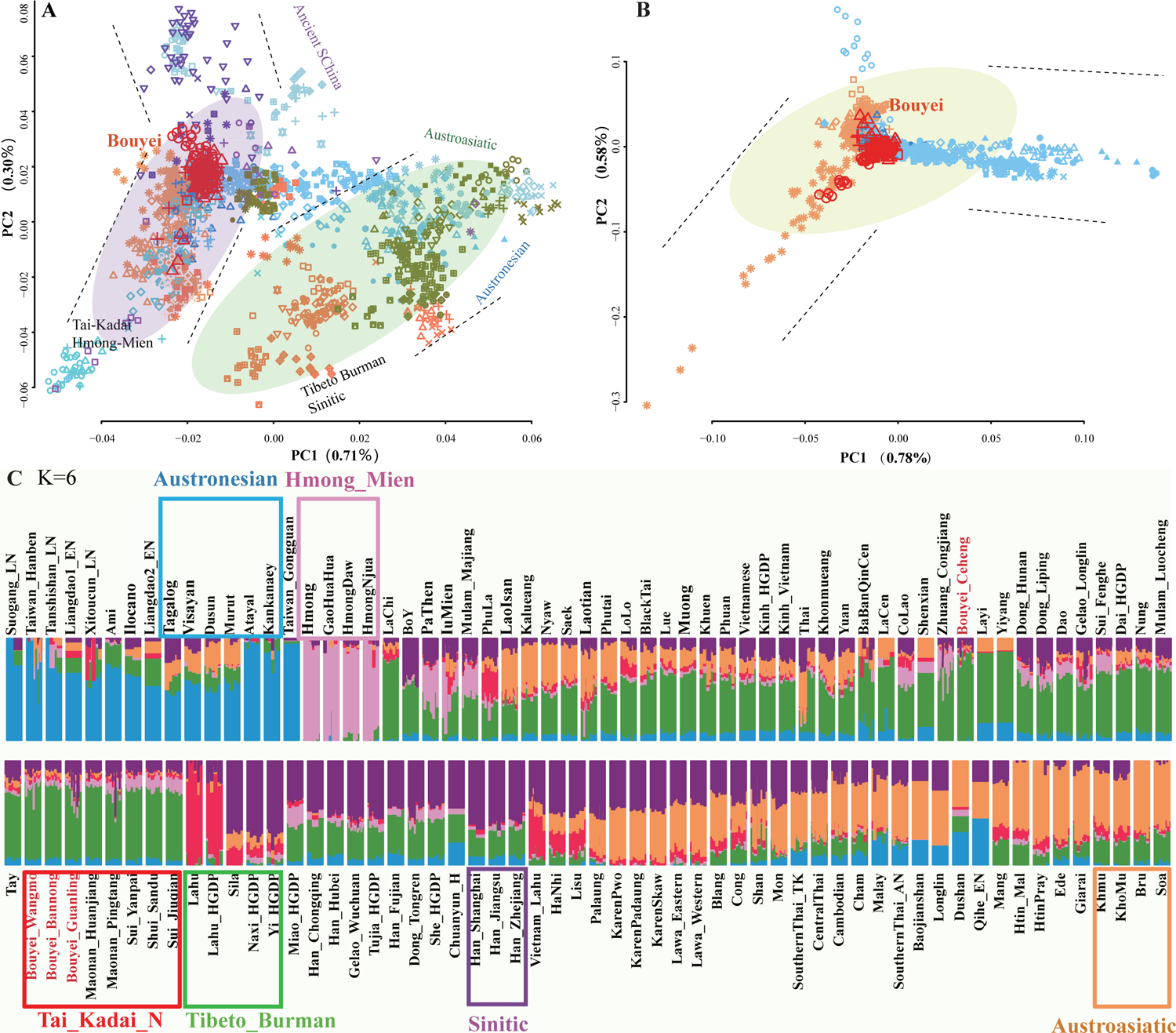
39 TK-speaking populations exist in the obvious genetic substructure. (**A**) 117 South Chinese population-based PCA was conducted based on the genetic variations of modern and ancient people projected onto it. (**B**) 39 TK-speaking population-based PCA was conducted based on the genetic variations of modern and ancient people projected onto it. (**C**) Population clustering patterns of both modern and ancient populations based on the “merged-H” dataset were revealed via model-based ADMIXTURE when six predefined ancestral sources were employed (K=6).

### Genetic substructure among 39 TK-speaking populations

The patterns of genetic affinity of TK-speaking populations inferred from PCA and model-based ADMIXTURE significantly differed from those observed in geographically distinct TB people from the Tibetan Plateau and other geographical edge populations. Most TK-speaking people had multiple ancestral components and were clustered in the intermediate position in the East Asian-scale PCA result. To further illuminate the genetic similarities and differences within geographically different TK-speaking populations and explore their fine-scale population substructure, we conducted the population genetic analysis and admixture modelling within 39 TK-speaking populations. The ADMIXTURE result revealed consistent patterns with the patterns observed in the PCA, which indicated that geographically different TK-speaking people possessed varying proportions of ancestral components and different genetic admixture histories. The increase of predefined ancestral sources (K values) from 6 to 10 generally led to a single ancestral population component being distinguished, suggesting their extensive genetic substructure and drift (**Figure S3**). To further investigate the genetic differentiation of TK-speaking populations, we constructed the phylogenetic tree based on 1-outgroup-*f_3_* among 117 Southern Chinese populations **(Table S3, Figure 3A**). We found that ancient people from South China, except for Gaohuahua, clustered more closely with modern AN speakers than other reference populations, and the 500-year-old Gaohuahua people clustered with HM-speaking people. We observed three main branches among modern populations. The upper branch included northern TK, northern TB, HM and their geographical neighbours. People from similar language families or geographical locations gathered with each other. The middle branch included southern TB and their neighbours; the lower branch included southern mainland TK and AN people. Bouyei groups generally clustered with geographically close TK-speaking populations, such as Maonan, Zhuang and Shui people from Guizhou province.

**Figure 3.**
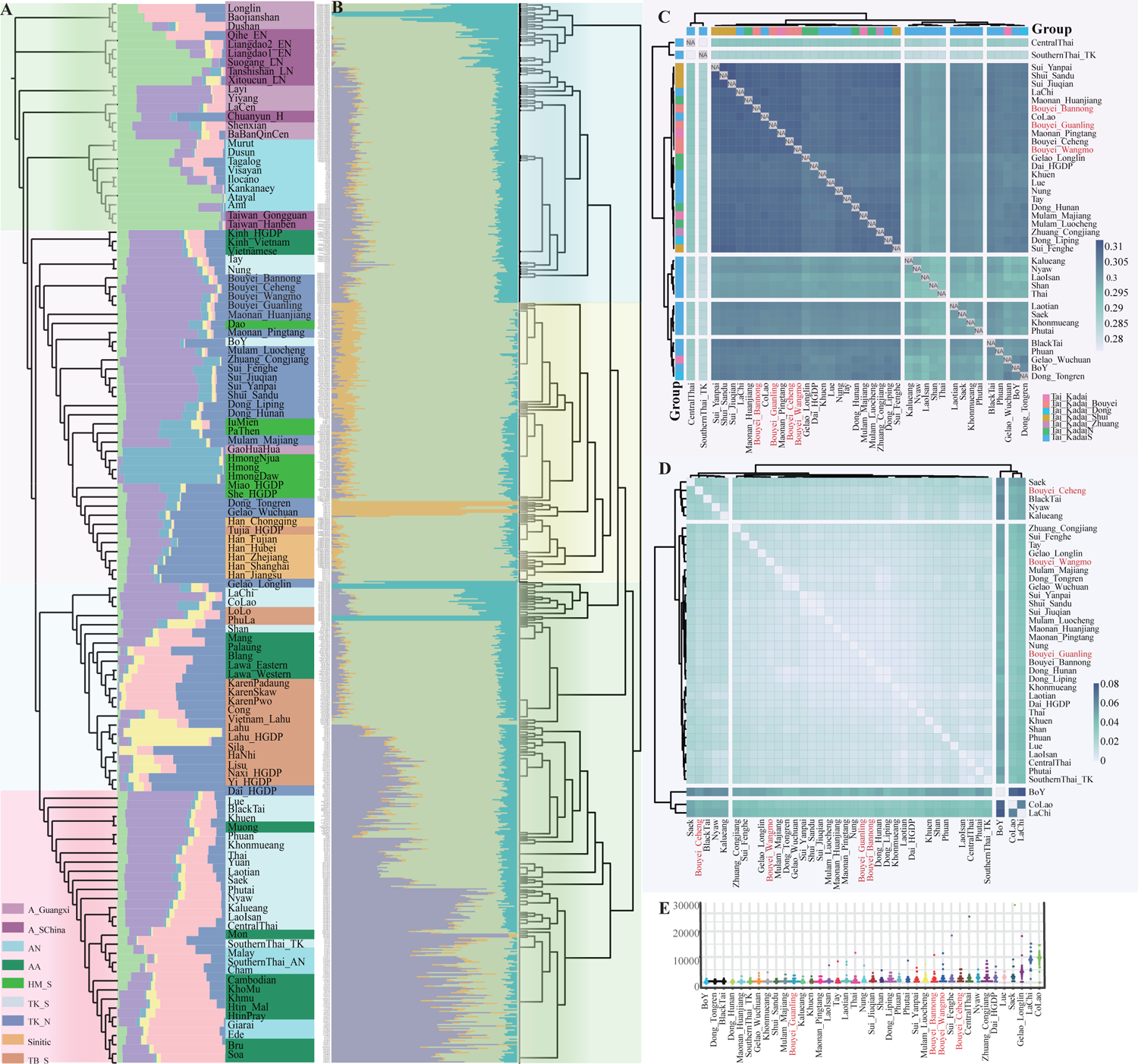
The genetic affinity of 39 TK-speaking populations. (**A**) A cluster topological tree based on the values of 1-outgroup *f*_3_ of 117 populations in Southern China and SEA. (**B**) The results of ADMIXTURE with K=4 and fineSTRUCTURE within 39 TK-speaking populations. (**C**) Heatmap of the shared genetic drift inferred from outgroup *f*_3_-statistics in the form *f*_3_(Studied Bouyei, TK; Mbuti), TK including 39 populations in Southern China and SEA. (**D**) Heatmap of the pairwise Fst genetic distances among 39 TK-speaking populations. (**E**) The mean lengths of runs of homozygosity for 39 TK-speaking populations.

We further explored the genetic differentiation and substructure using haplotype-based fineSTRUCTURE and admixture model reconstructions based on the allele frequency spectrum among 796 individuals (**Figure 3B**). Haplotype-based clustering patterns were consistent with the model-based ADMIXTURE ancestral composition, in which most TK individuals possessed primary simulated ancestry maximised in the northern Guizhou and southern Thailand people. The pattern of genetic similarity inferred from the 1-outgroup-*f*_3_ heatmaps also revealed differentiated patterns of shared alleles between northern and southern TK-speaking populations (**Figures 3C and S4**). We observed a stronger genetic affinity between Bouyei and geographically proximate Chinese TK-speaking populations, consistent with the identified genetic relationship based on the pairwise F_ST_ genetic distance (**Figure 3D, Table S4)**. Moreover, we investigated the correlations between the length and quantity of the runs of homozygosity (ROH) across various ethnic minorities and further explored the distribution pattern (**Figure 3E**). The total lengths of ROH in TK-speaking populations from MSEA were comparatively longer, whereas Chinese TK-speaking populations exhibited a similar distribution pattern. Generally, we highlighted geography-related genetic substructure within 39 TK-speaking populations, including Southern TK-speaking populations in MSEA and Northern TK-speaking populations in Southern China.

**Figure 4.**
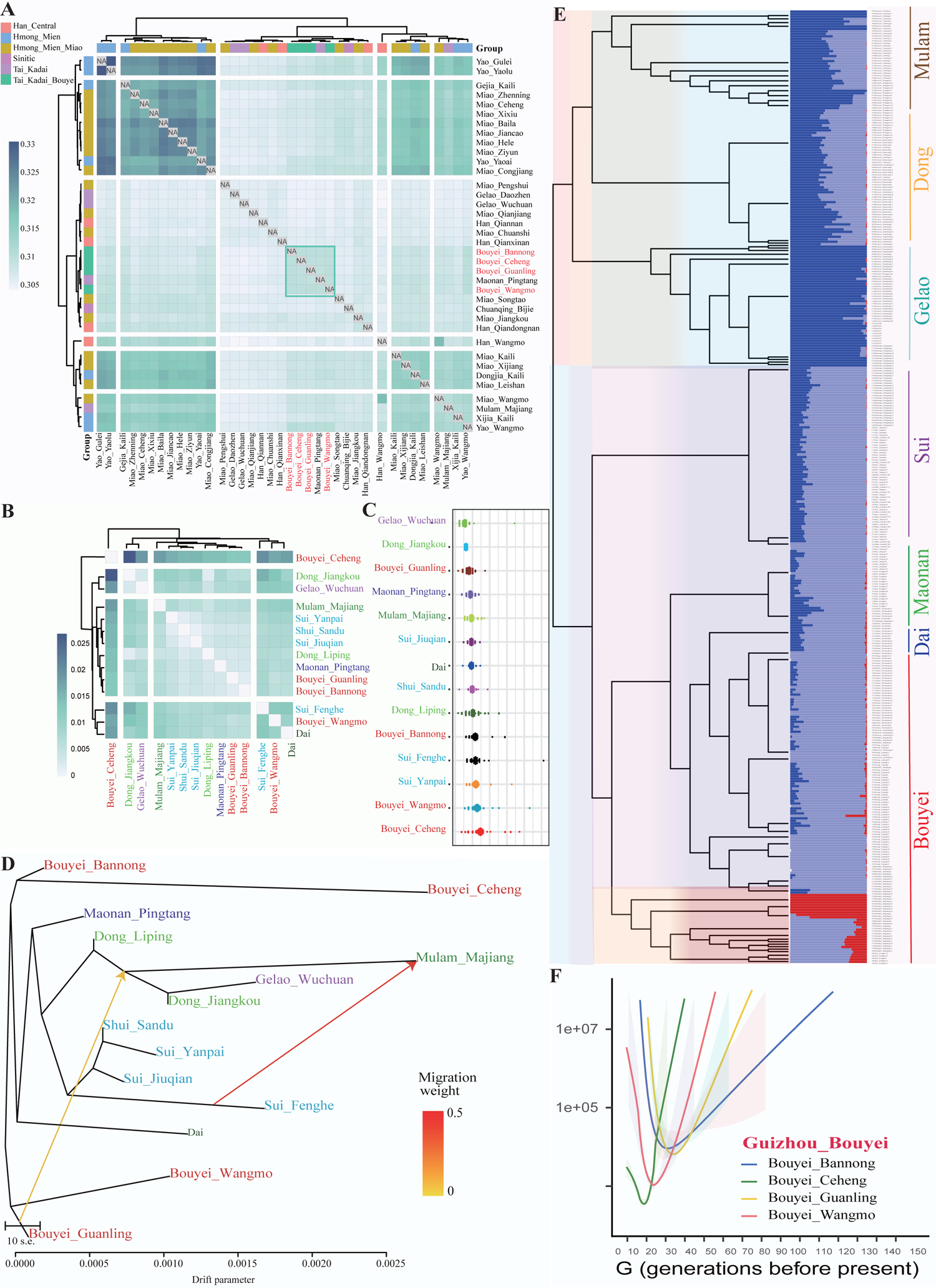
Four Bouyei populations have a close genetic relationship with surrounding populations in Guizhou. (**A**) Heatmap of the shared genetic drift inferred from outgroup *f*_3_-statistics in the form *f*_3_(Studied Bouyei, Reference; Mbuti). (**B**) Heatmap of the pairwise Fst genetic distances among 14 TK-speaking populations in Guizhou. (**C**) The mean lengths of runs of homozygosity within 14 TK-speaking populations. (**D**) The phylogenetic relationship showed the close genetic affinity between Bouyei and other TK-speaking populations in Guizhou. (**E**) The results of ADMIXTURE with K=3 and fineSTRUCTURE within 14 TK-speaking populations in Guizhou. (**F**) The effective population size of four Bouyei from 150 generations before the present.

Population genetic structure analyses in the context of East Asians and regional TK-speaking people identified one unique ancestry dominant in Guizhou TK-speaking people. The complexity of geographical environments and unique cultures indirectly provided favourable conditions for forming various ethnic groups and genetically differentiated population structure(23). To deeply illustrate the confidence of the identified new ancestry component and explore their interaction with geographically different Guizhou people, we performed a population comparison analysis among Guizhou populations, including HM/TB/Sinitic and TK-related populations. We observed that Bouyei shared the most alleles with surrounding TK-speaking populations and other geographic proximity ethnic minorities (**Figure 4A**). Shared alleles and haplotypes further confirmed the high genetic affinity among geographically different TK-speaking populations and interacted with each other frequently (**Figures 4B and S5**). The ROH values of Bouyei populations were comparatively higher, and the distribution patterns showed remarkable similarities to neighbouring ethnic minorities such as Dong and Sui (**Figure 4C**), which represented the possibility of inbreeding(28). We also reconstructed the maxim-likelihood-based TreeMix among 14 TK-speaking populations to investigate the phylogenetic relationship (**Figure 4D**). TreeMix-based phylogenetic relationship demonstrated the population substructure among TK-speaking populations and indicated that the BBN had close genetic affinities with other geographically different Bouyei groups when two gene flow events occurred. The effective population size (N_e_) within the past 150 generations inferred the bottleneck events at different times and placed genetic diversity among TK-speaking populations (**Figure 4F**). Additionally, we explored the genetic differentiation and the patterns of the fine-scale genetic structure using fineSTRUCTURE (**Figure 4E**), which was consistent with the patterns of population substructure revealed by the pairwise coincidence matrix (**Figure S6**).

**Figure 5.**
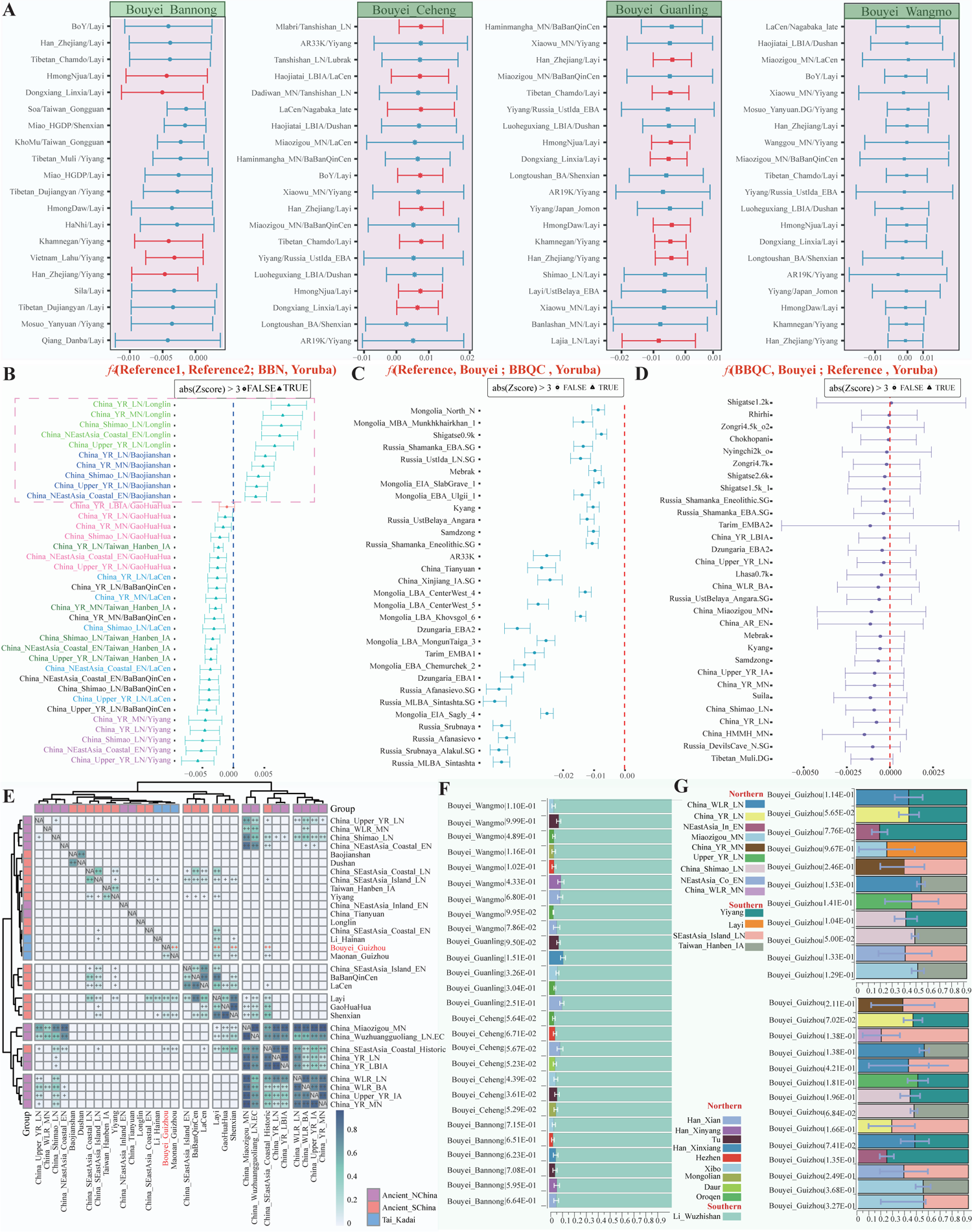
Admixture events and demographic history. (**A**) admixture-*f*_3_-statistics in the form *f*_3_(Modern /Ancient Reference 1, Modern /Ancient Reference 2; Bouyei). Lowered the threshold and set the Z-score lower than −2 and visualise the top highest 20 values. Red represents Z-values that are significant, and blue represents Z-values that are not significant. (**B**) *f*_4_-statistics test in the form of *f*_4_(Reference population1, Reference population2; BBN, Mbuti) based on middle-density merged 1240K dataset to identify the genetic affinity between BBN and other references. (**C**) *f*_4_-statistics in the form of *f*_4_(Reference, Meta-Bouyei; BBQC, Mbuti) to explore the genetic association with Meta-Bouyei and BBQC. (**D**) *f*_4_-statistics in the form of *f*_4_(BBQC, Meta-Bouyei; Reference, Mbuti) to verify whether the Meta-Bouyei were influenced by other reference except BBQC. (**E**) Pairwise qpWave results showed the genetic homogeneity and heterogeneity between Meta-Bouyei and other ancient populations. (**F**) Two-way admixture models showed that both modern Northern and Southern populations contributed to the formation of four Bouyei people. The error bar indicates the stand errors of predicted proportions of ancestors obtained from qpAdm. (**G**) Two-way admixture models showed that both ancient Northern and Southern populations contributed to the formation of the Meta-Bouyei people. The error bar indicated the stand errors of predicted proportions of ancestors obtained from qpAdm.

### The shared alleles revealed by F-statistics

To explore the detailed demographic history and possible ancestral sources of Bouyei populations, we conducted admixture *f*_3_-statistics in *f*_3_(Reference population1, Reference population2; Studied populations) based on the HO dataset (**Figure 5A**). We did not observe significant *f*_3_ values in the Bouyei groups. We cannot completely exclude the ancient or recent admixture events with the following strong population bottlenecks. We further performed ChromoPainterv2 to paint the ancestral haplotype composition and ran a fastGLOBETROTTER based on the haplotype sharing to identify ancestral sources, date and describe admixture events of our targeted populations. For BBN groups, the best-guess conclusion for admixture was “unclear signal”, which provided evidence about the unique population demographic history and suggested its relatively isolated genetic background. We then performed a series of *f*_4_-statistics to explore genetic differentiation and gene flow events between Bouyei and other populations. Interestingly, we did not observe significant negative signals except spurious signals (**Figure S7**). Meanwhile, we observed the same result based on the genetic variations of modern populations (**Table S5**), which confirmed relative genetic homogeneity within four Bouyei groups.

To further explore the genetic affinity between BBN and reference populations, we conducted *f*_4_-statistics in the form *f*_4_(Reference population1, Reference population2; BBN, Mbuti). When focused on ancient populations, we found that studied groups were affected by Northern and Southern gene flow. BBN shared more alleles with ancient Northern East Asian populations than with early Neolithic Southern groups (Baojianshan/Longlin). BBN received more genetic influence from ancestry related to Guangxi historical people (BBQC), earlier Fujian Neolithic individuals and Taiwan Island Hanben people compared with ancient YRB farmers (**Figure 5B**). For modern East Asian populations, BBN has a closer genetic relationship with Southern populations than with Northern Chinese (**Table S6∼9**). Within Southern Chinese populations, we observed that Bouyei populations shared more alleles with HM in *f*_4_ (AN, HM; BBN, Mbuti). Consistent with the genetic relationship inferred from outgroup *f*_3_ (**Figure 4A**), Bouyei populations had a closer genetic affinity with other geographically close TK-speaking populations in *f*_4_ (TK, HM; BBN, Mbuti). Besides, to further explore general ancestral sources and improve the statistical power of Bouyei populations, we merged four Bouyei groups into a meta-Bouyei population as their genetic homogeneity identified via pairwise qpWave analysis (**Figures S7∼8**). According to archaeological evidence(29), BBQC was a historical population living in Guangxi Province 1500 years ago, which was considered the direct ancestor of the TK-speaking people. We performed *f*_4_-statistics in the form of *f*_4_(Reference populations, Meta-Bouyei; BBQC, Mbuti) and found abundant significant negative *f*_4_ values, which suggested that our studied groups shared more alleles with BBQC compared with other references (**Figure 5C**). Additionally, Bouyei populations were influenced by gene flows from Tibetan-related ancestral sources (Muli Tibetan: Z= −2.572) in the form of *f*_4_(BBQC, Meta-Bouyei; Reference populations, Mbuti) (**Figure 5D**), which was consistent with the heterogeneity observed within Bouyei populations and BBQC (**Figure 5E**). Overall, different types of *f*_4_ analyses shed light on a strong genetic affinity within TK-speaking populations and demonstrated a common origin in the Bouyei population.

We build two-way qpAdm admixture models with the Han and other Altaic people as the Northern Chinese surrogates and the island Li people as the Southern surrogate to directly explore the admixture sources and proportions of our newly-studied Bouyei people. We observed that the contribution from Li-related ancestral sources ranged from 0.8860 to 0.9690. Northern ancestry sources just occupied a low proportion, which supported that Bouyei people originated from Southern Chinese indigenous people and were affected by gene flow events from Northern populations (**Figure 5F**). Three-way qpAdm models also provided a good fit for Bouyei’s admixture history **(Figure S9)**. Quantitative *f*-statistics demonstrated that Bouyei had a closer genetic relationship with Guangxi’s historical populations (**Figure 5C∼D)**. Therefore, we used ancient Northern YRB farmers and Southern rice farmers as surrogates and confirmed our hypothesis that Bouyei populations could be modelled as a north-south ancestral admixture (**Figure 5G**). To further pinpoint the date of admixture events between Northern and Southern populations in a wide range of time, we used the ALDER (Admixture-induced Linkage Disequilibrium for Evolutionary Relationship) and discovered complex genetic admixture in Bouyei populations **(Table S10)**. We found admixture occurred in BBN about 55.86±26.51 generations (1536±742 CE) in the Han-Yao model and 94.37±35.94 generations (2614±1006 CE) in the Han-Dongjia model.

### Natural selection signatures and biological adaptation

Our identified complex genetic admixture processes and environmental selection forces may contribute to TK populations’ unique landscape of the biological adaptative variants or genes. We used multidimensional techniques based on the allele frequency spectrum and haplotype homozygosity to comprehensively characterise the biological adaptative features of our firstly-reported TK lineage in Guizhou. First, we applied population branch statistics (PBS) with Northern Han Chinese as the ingroup reference population and the merged European from HGDP genomic resources as the outgroup to identify the putatively adaptative signatures that occurred in Bouyei people after separating from Han Chinese. Since the substantial homogeneity in Bouyei people, we combined four Bouyei groups into the merged meta-Bouyei as the targeted populations. Loci with the PBS values over the 99.99th percentile were regarded as the candidate selection variants. We identified 141 PBS-identified adaptative genes on chromosomes 1, 2, 3, 6, and 11 (**Figure S10, Table S11**). We performed functional enrichment analysis based on the PBS-based signatures (**Figure 6A**). Enrichment results showed the selection-related genes were associated with lipid metabolisms [regulation of Linoleic acid (LA) metabolism (R-HSA-2046105)], immune [Intestinal immune network for IgA production (hsa04672)], nervous development [regulation of nervous system development (GO:0051960) and regulation of neuron projection development (GO:0010975)] and other development and proliferation pathways [multicellular organismal homeostasis (GO:0048871), G protein signalling pathways (WP35), cellular response to UV-B (GO:0071493), and adenylate cyclase-activating G protein-coupled receptor signalling pathway (GO:0007189)].

**Figure 6.**
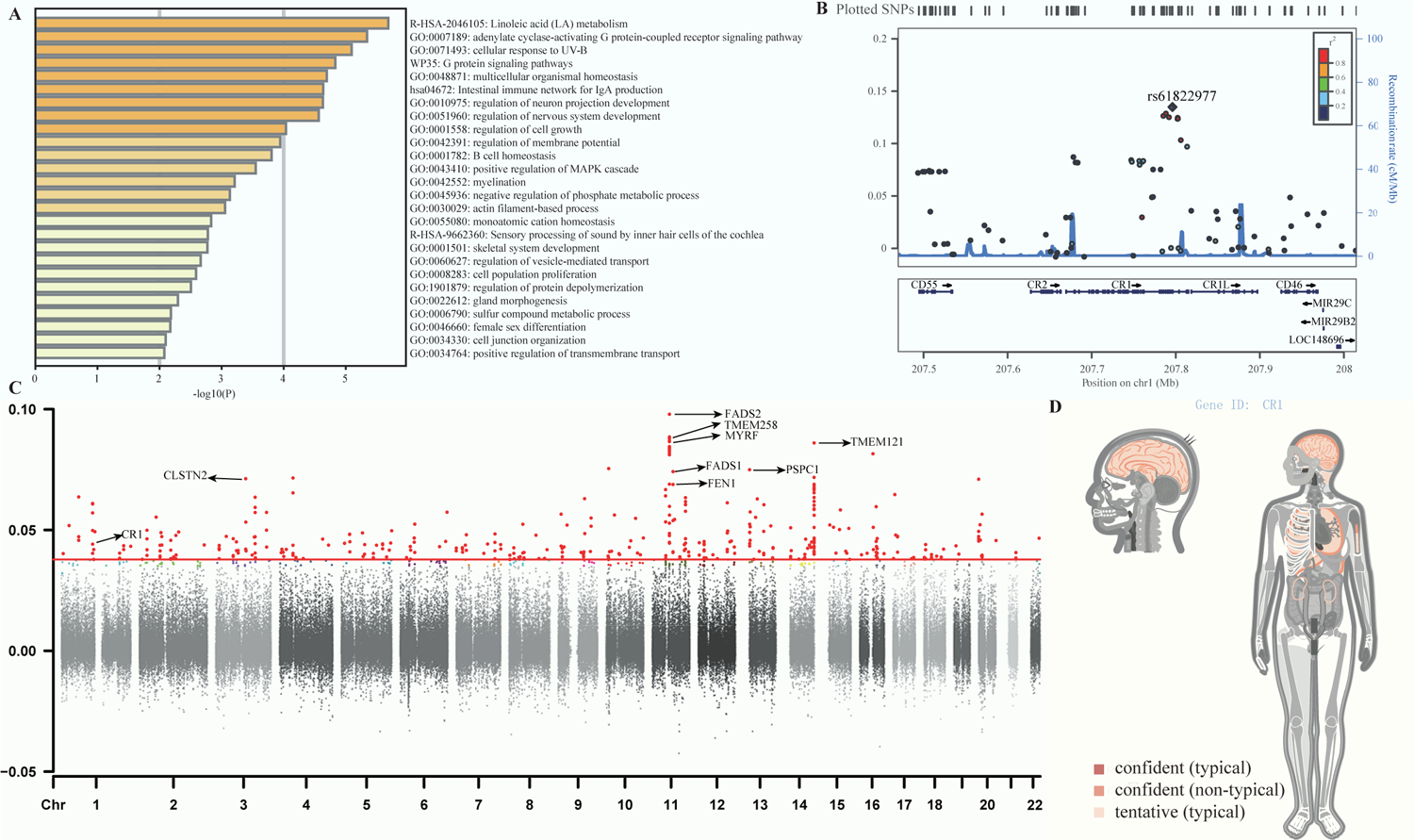
Positive natural selection signals and other relevant variations in Bouyei populations. **(A)** Functional enrichment for the candidate genes in which the PBS values were top 100 in the form of (Meta-Bouyei-Shaanxi_Han-European) according to Metascape online software. The enrichment p values showed genes associated with the pathway. (**B**) The concerning adaptive variant (rs61229077) which was located in the *CR1* gene on chromosome 1 was marked, and other SNPs were coloured based on the pairwise linkage disequilibrium with the target variant. (**C**) Manhattan plot showing the PBS values in genome-wide scanned for the Bouyei population in Guizhou, using the Miao_GZ and Shaanxi_Han as reference populations. The 99.9th percentiles of the PBS distribution were shown as red lines. PBS values over the 99.9th percentile were marked in red, and PBS values under the 99.9th percentile were coloured as dark dots. Elsewise, some of the genes were labelled with its name. (**D**) The *CR1* gene affected relevant organs, systems, regions, and germ layers, according to the Gene organiser.

The highly noteworthy natural selection signals for Bouyei were identified in genes related to lipid metabolisms and physical features, such as *FADS1* (Fatty Acid Desaturase1), *FADS2* (Fatty Acid Desaturase2) and *EDAR* (Ectodysplasin A Receptor). The strongest PBS selection signal was identified in *EDAR* (PBS=0.3180) located on chromosome 2 (rs922452), which is a member of the tumour necrosis factor receptor family and played a predominant role in hair straightness, skin tone, facial features flatness and sweat gland density in East Asians(30). *FADS* is associated with lipid metabolism determined the long-chain polyunsaturated fatty acid (PUFA) level, mainly including genes: *FADS1* and *FADS2*. *FADS* encodes the fatty acid desaturase enzymes, which regulates the unsaturation of fatty acids by translating the short-chain to long-chain PUFA (28). *FADS*-rs174570 had the top selection signal (PBS=0.2533). Previous genetic studies have reported that *FADS* levels were associated with Lipid Metabolism Disorder and Hypobetalipoproteinemia(31) (32). Ancient Southern Chinese people experienced a long history of infectious diseases. Recent modern and ancient European genomes have demonstrated that infectious disease risk increased since the early Neolithic period, and autoimmune-related disease risk and adaptative evolution occurred in the post-Bronze Age(33). We also identified immune-related putatively adaptative variants, multiple variants observed in *HLA-DPA1* (Major Histocompatibility Complex, Class II, DP Alpha 1) and *CR1* (Complement C3b/C4b Receptor 1, Knops Blood Group). According to polypeptides produced by extracellular proteins, *HLA-DPA1,* as a protein-coding gene, mainly plays a crucial role in the immune system. The key mutations in *HLA-DPA1* are associated with the genetic susceptibility of rare autoimmune diseases, such as Granulomatosis with Polyangiitis(31). Epidemiological data documented the high incidence of Malaria in South China, and Southern Chinese people possessed the adaptive features during persistent pathogen exposure. *CR1* plays a vital role in the mutual effect within the P. falciparum and interrelated hosts at different levels. Genetic variations in *CR1* lead to *CR1* deficiency, which happens in regions with high incidences of Malaria, yet this mechanism can prevent severe Malaria. In detail, parasitised red blood cells (RBCs) invaded by severe Malaria typically adhere to complement receptor one located on other uninfected RBCs and play the corresponding physiological roles in forming clumps of cells that are also referred to as rosettes (31, 34–36). In addition, we observed that the rs61822977 with the highest PBS value on chromosome 11 showed a robust biological adaptation and association among other identified *CR1* related-SNPs. Besides, we further drew the regional plot for the top-ranked SNP rs61822977 and other nearby variants in 200kb based on linkage disequilibrium. Other immune-related putatively adaptative variants (*CD55*, *CR2*, *CR1L* and *CD46*) were also marked (**Figure 6B**). However, because of the complexity of the host-parasite interactions, more research was needed to elucidate the vital molecule makers and corresponding biomedical mechanisms acting on the pathogenesis of Malaria.

To recognise the differentiated regional-specific natural adaptation signals of the Bouyei people, we also calculated PBS using neighbouring Miao and Northern Han as the ingroup and outgroup. We identified 132 adaptative genes within the candidate selection variants over the 99.99th percentile in the PBS values (**Figure 6C, Table S12)**. Except for genes like *FADS1*, *FADS2* and *CR1* above, we also found strong candidate genes related to familiar diseases. For example, *CLSTN2* (Calsyntenin 2) lies on chromosome 3 and is relevant to Astigmatism; *TMEM258* (Transmembrane Protein 258) level is related to Spinocerebellar Ataxia; another gene *TMEM121*(Transmembrane Protein 121) plays an essential role in the component of membrane; *FEN1*(Flap Structure-Specific Endonuclease 1) is linked to Xeroderma Pigmentosum through the base excision repair(37). *MYRF* (Myelin Regulatory Factor) lies in chromosome 11 and encodes a transcription factor essential in developing the myelin central nervous system and myelination. It can be regarded as increasing gene expression(38) and further influencing myelin production, while others directly facilitated myelin gene expression. In addition, for the *CR1* gene, which also was identified in regional-specific analyses, we performed Gene ORGANizer (http://geneorganizer.huji.ac.il/) to explore the association between the *CR1* gene and organ systems. We observed that our identified putative adaptative genes were potentially linked and further influenced by our brain and lung, corresponding to the physiological mechanism related to the Malarial pathogens that invaded and damaged the human body (**Figure 6D**). Using the enrichment database in Metascap, we found these candidates were related to essential physiological functions of the human body, such as cell-cell adhesion, neuronal system, development process, and immune system process. Besides, we performed pathway and process functional enrichment with the three ontology sources: GO Biological Processes, Reactome Gene Sets, and WikiPathway, to investigate the detailed functional connections. Within 132 candidates under natural selection, we listed 20 functional groups in total (**Figure 7A, Table S13)**. The functional category was considered significant when − log10_P value_ > 2. Cell morphogenesis involved in neuron differentiation (GO:0048667) and regulation of proteolysis (GO:0030162) accounted for the highest proportion, approximately 10.08%, respectively. We also calculated the five PBS values using different ethnic minorities in Guizhou, including Zhuang/Shui/Mulam/Maonan and Gelao as the ingroup and Northern Han as the outgroup, respectively (**Figure 7B**) to detect the specific signals of adaptive evolution among Bouyei people populations. Except for two genes (*CLSTN2* and *CR1*) mentioned above, *PTPRD* (Protein Tyrosine Phosphatase Receptor Type D) gene was identified in all six analyses, which was a member of the protein tyrosine phosphatase (PTP) family and relevant to Restless Legs Syndrome and Chromosome 9P Deletion Syndrome. In six analyses of different ethnic groups in Guizhou, the *CR1* gene was screened simultaneously, verifying that *CR1* played a vital role in resistance to Malaria in Guizhou. Therefore, based on the 10K Chinese People Genomic Diversity Project (10K_CPGDP) database, we explored the distribution of allele frequency about *CR1-*related mutation(rs7542544) in diverse spatial (**Figure 7C**) and focused on the frequency distribution in China and especially in Southern China (**Figure 7D**). We observed that this mutation (rs7542544) presented an intense natural selection signal, and its derived haplotype was highly expressed in Southern China, which also conformed to the frequency distribution in Sardinia, Papua New Guinea, Bantu_Kenya and other Malaria-endemic regions(30, 34, 36). It was worth mentioning that this phenomenon about high frequencies of the derived allele also occurred in TK-related populations with different geographical environments, which further indicated that their common ancestors were affected by strong natural selection and followed by the occurrence of population admixture, migration as well as genetic drift and bottleneck events. A typical example was the frequency distribution pattern in Hainan Li, which can be representative of the coastal TK-speaking population(16). Furthermore, other eight loci also confirmed Malaria prevalence among TK-speaking populations **(Table S14)**. Consequently, RBC *CR1* deficiency caused by high expression of *CR1*-derived alleles was profoundly common in Southern China malaria-endemic regions such as Guizhou and other SEA. The polymorphisms with *CR1* deficiency conferred protection against severe Malaria(31, 34–36).

**Figure 7.**
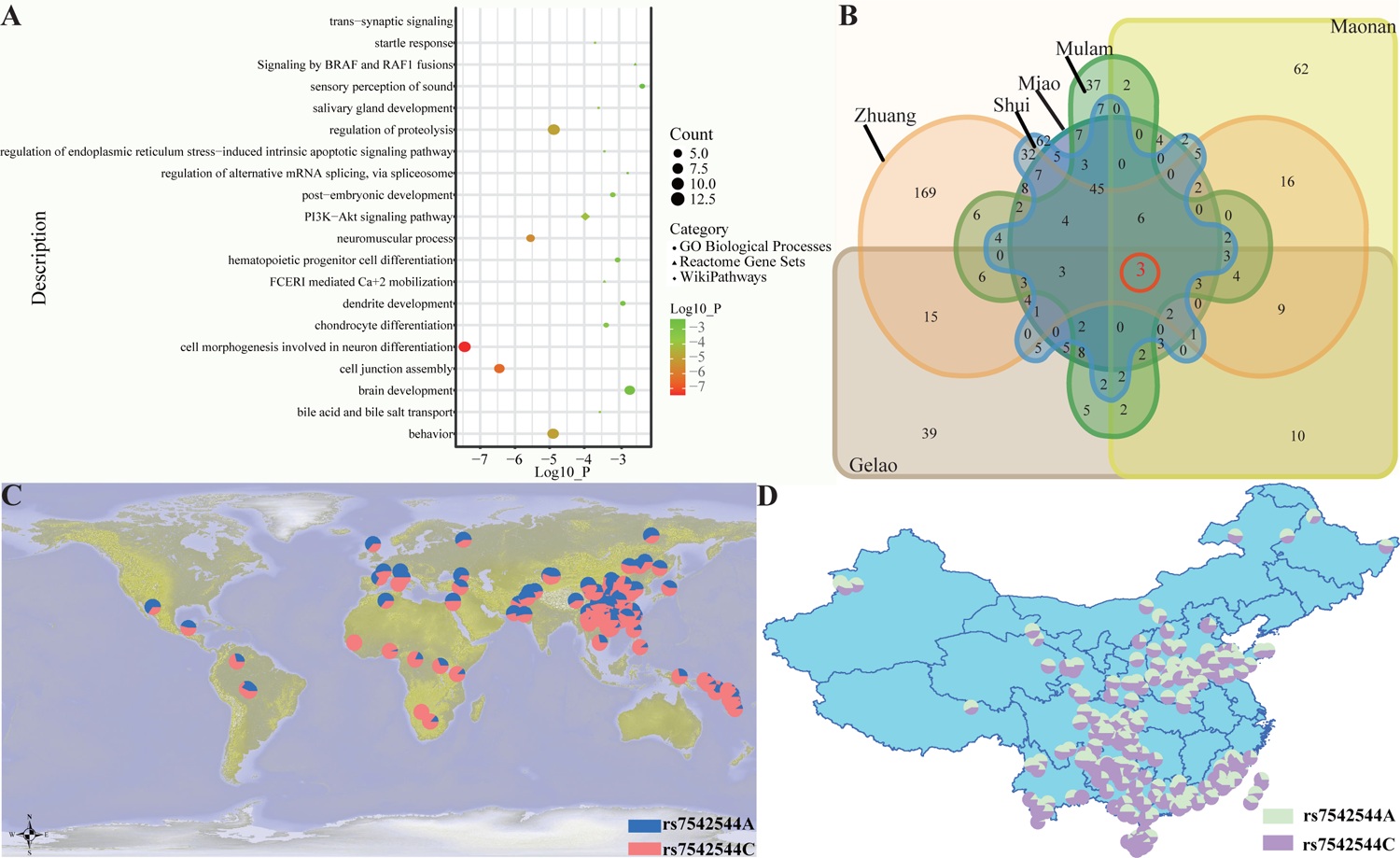
Regional-specific natural selection signals. (**A**) Functional enrichment for the candidate genes in which the PBS values were over 99.9th percentiles in the form of (Meta-Bouyei-Miao_GZ-Shaanxi_Han) according to Metascape online software. Also, these candidates formed the top 20 clusters based on three categories; “Coun” is the number of genes in these candidates for each cluster; “Log10” was the p-value in log base 10, when − log10_(P value)_>2 it indicated the functional category was considered significant. (**B**) Venn plot represented the overlaps of positive natural candidates in six group-PBS in the form of (Meta-Bouyei-Miao/Gelao/Zhuang/Shui/Mulam/Maonan-Shaanxi_Han). (**C**) The frequency distribution of *CR1* (rs7542544 A/C) was associated with the genetic variant based on the 10K_CPGDP global database. (**D**) The frequency distribution of *CR1* (rs7542544 A/C) was associated with the genetic variant based on the 10K_CPGDP database in China.

## Discussion

### Demographic history and population structure within TK-speaking populations

Rice farming originated in the Yangtze River and led to the development of the Southern Chinese TK/HM/AA/AN-related populations, which occupied an essential role in population migration southward to SEA(8, 38, 39). The ancient “Baiyue” living in Southeast China was considered to be the ancestors of the present-day TK-speaking populations. The TK-speaking populations were the indigenous people of Southern China. Continuous population admixture and isolation events contributed to their specific genetic structure and evolutionary history. Previously, fewer studies had spotlighted inland and integrated groups and more research paid attention to the genetic structure of the single and coastal groups. One of the typical representatives was Hainan Li, which had a relatively isolated genetic background and complex evolutionary history(13, 37). Therefore, the genetic origin and phylogenetic relationship within TK-speaking populations remained to be further characterized(32, 40–42). We collected and genotyped 24 Bouyei individuals from Guizhou Bannong and merged newly-generated data with previously published data to produce three datasets: low-density merged HO dataset(26, 27); middle-density merged 1240K dataset, and high-density merged WGS dataset(43–49) to provide new insights into the demographic history, population structure and human genetic diversity of TK-speaking. The ADMIXTURE result based on a high-density dataset identified a unique genetic component maximised in TK-speaking Bouyei, indicating that Bouyei populations possessed a unique genetic structure (**Figure 1A**). As the high-density dataset lacked SNP data on other TK-speaking populations from MSEA, we further included populations genotyped via the Affymetrix Human Origin array and explored the general patterns of genetic structure among 39 TK-speaking populations and their relationship with other ancient/modern humans (**Figures 1B∼2**). Our results showed that TK-speaking populations from MSEA and China showed similar genetic structures and close genetic affinity (**Figures 2B∼C, 3**). Whereas we identified population genetic substructure related to their geographic distribution patterns. 39 TK-speaking populations could be roughly divided into two clusters: Southern TK-speaking populations from MSEA and Northern TK-speaking populations from China, which showed apparent genetic differentiation. Bouyei groups and surrounding TK-speaking populations (Maonan) clustered together and showed more allele sharing. TK-speaking populations from MSEA showed similar genetic structures and relationships.

Guizhou Province is located in the Yungui Plateau and has mountainous topographical features, resulting in an isolated geographical environment and a unique genetic landscape(23). Concentrating on 14 populations in areas with high ethnic and linguistic diversity based on a high-density dataset, we identified their high genetic affinity and similar genetic origins (**Figure 4**). We performed admixture *f*_3_ and GLOBETROTTER to detect the admixture signatures of the BBN population (**Figure 5A**). Both genetic alleles and haplotype blocks confirmed that Bouyei has a unique evolutionary history. Moreover, a high degree of genetic homogeneity within four Bouyei groups and widespread heterozygosity within TK-speaking populations were demonstrated through the observed results of *f*_4_(Studied1, Studied2; Reference, Mbuti) and qpWave (**Figure S8∼9**). In addition, a previous study illustrated that 1500-year-old BBQC people in Guangxi were reported as the direct ancestral source of the modern TK people. We observed that the estimates of *f*_4_(BBQC, Bouyei; Reference populations, Mbuti) were consistent with the previous study. Bouyei populations shared more alleles with BBQC and were influenced by gene flow events from neighbouring minorities (**Figures 5C∼D**). The results of *f*_4_(Reference population1, Reference population2; Bouyei, Mbuti) also revealed widespread gene flow events from surrounding ethnic groups (**Figure 5B**). Finally, the qpAdm-based two/three-way admixture models and ALDER further confirmed their potential admixture proportions and times (**Figure 5F∼G**). Bouyei populations originated from southern indigenous people and were affected by gene flow events from northern populations.

### Local adaptation of Guizhou populations

To understand the genetic basis of local adaptation signatures in Bouyei populations, we performed PBS analyses with Northern Han Chinese as the ingroup and the European_HGDP as the outgroup and identified 141 adaptative genes with the values over the 99.99th percentile (**Table S8**). According to the functional enrichment analysis, we observed different signalling pathways, including immune, lipid metabolism, physical traits, and so on (**Figure 6A**). *HLA-DPA1* mainly plays a central role in the immune system by presenting peptides derived from extracellular proteins(31). *FADS1* and *FADS2* mainly encode the fatty acid desaturase enzymes and control the unsaturation of fatty acids(28). The *EDAR* gene is associated with typical characteristics in East Asian features, including the shovel shape of upper incisors, hair straightness, and facial characteristic(50–53). The physiological functions associated with *CR1* mainly include opsonisation, control of complement activation, and removal of immune complexes (ICs). *CR1* deficiency was also related to Malaria resistance, which can greatly reduce the resetting of the rosette. Specifically, RBCs invaded by severe Malaria typically adhered to complement receptor one located on other uninfected RBCs and played the corresponding physiological roles in a rosette formation, causing the severe obstruction of the cerebral microvasculature thus resulting in pathologic changes in cerebral malaria(31, 34–36). Furthermore, we performed the regional plot for the top-ranked SNP rs61822977 and found nearby immune-related putative adaptive variants (*CD55*, *CR2*, *CR1L,* and *CD46*) in 200kb based on linkage disequilibrium (**Figure 6B**). We also calculated PBS using the neighbouring Miao and northern Han as the ingroup and outgroup to explore the differentiated regional-specific natural adaptation signals within the Bouyei people. We identified 132 adaptative genes within the candidate selection variants over the 99.99th percentile in total (**Figure 6C, Table S9**), which were related to essential physiological functions of the human body through the enrichment database in Metascap (**Figure 7A, Table S10**). To further detect the specific signals of natural selection within the Bouyei populations, we used Zhuang/Shui/Mulam/Maonan and Gelao in Guizhou as the ingroup and northern Han as the outgroup to calculate the other five PBS values (**Figure 7B**). Interestingly, the *CR1* gene was screened simultaneously, suggesting that *CR1* played a vital role in resistance to Malaria in Guizhou. Besides, we found other strong candidate genes, such as *CLSTN2, TMEM258, TMEM121, FEN1,* and *MYRF*, which plays different roles in relevant organs and systems, except for genes such as *FADS1*, *FADS2,* and *CR1* above. For example, *MYRF* is a well-known gene located on chromosome 1 and encodes a transcription factor that is required for central nervous system myelination(54). Combined with the frequency distribution of nine related loci based on the 10K_CPGDP database, we found the derived haplotype of *CR1* was strongly selected and highly expressed in Southern China such as Guizhou and MSEA, resulting in Malaria resistance in endemic areas (**Figure 7C∼D**)(31, 34–36).

Guizhou has been described as a “miasmatic region”, with endemic diseases such as Malaria epidemics that seriously affected the local population’s health and further influenced social and economic development(55). Before liberation, the annual number of Malaria cases in Guizhou Province was between 2 million and 3 million, representing 25-30% of the total population. For decades, the government has vigorously pursued a comprehensive prevention and control strategy focusing on controlling the source of infection. However, with the increasingly frequent global economic, a mass of travel, and business affairs abroad from Malaria-endemic countries and regions, the imported Malaria threat will continue to exist in Guizhou province. The formation of IC was a prominent feature of Malaria infection, which can generate proinflammatory cytokines, thus stimulating macrophages and monocytes, and *CR1* also played a crucial role in IC clearance due to its high affinity for C3b and C4. Erythrocyte *CR1* (*E*-*CR1*) binds the ICs in the peripheral blood through “immune adherence”, which can transport them to the phagocytes in the liver or spleen to further remove them from the circulation. Other research also showed that erythrocytes with high levels of *CR1* would carry a further quantity of ICs, which stimulated the production of proinflammatory cytokines, thus increasing the incidence of cerebral Malaria augmenting. Therefore, the *CR1* levels on erythrocytes and relevant polymorphisms had been associated with the response to falciparum Malaria in Malaria-endemic regions(31, 34–36). Meanwhile, the severity of falciparum Malaria has been linked to several human genetic factors such as including sickle-cell disease, thalassemia, and G6PD deficiency. These diseases via erythrocytic defects were considered as the selective pressure behind(55).

## Conclusion

We merged newly generated and publically available genome-wide SNP data and formed one aggregated dataset to explore the population structure, evolutionary history and biological adaptation within 39 geographically different TK-speaking populations. We found that Bouyei groups were genetically similar to neighbouring TK-speaking populations. Furthermore, due to its unique genetic structure, Bouyei-related ancestry can be used as an optimal representative of inland TK-related populations. The shared haplotypes and alleles showed genetic heterozygosity among 39 TK-speaking people from diverse geographical environments. In addition, we found that 39 TK-speaking populations exhibited the prominent geography-related population substructure, and the clustering patterns were associated with their extensive differences in genetic diversity. We found that four Bouyei people had strong genetic homogeneity, simulated admixture time and constructed admixture models within Bouyei populations, further providing evidence for their distinct genetic origins. We also identified north-to-south admixture events which were consistent with genetic affinity and historical population movements. It was confirmed that Bouyei could be modelled as an admixture of minor Northern Chinese populations and major Southern Chinese populations, and ancient South Chinese people constituted the majority of Bouyei ancestry. We identified several candidates through the population and regional-specific analyses, including the *CR1* gene associated with immunity and Malaria resistance and other genes with metabolic evolution.

## Materials and methods

### Sample collection and DNA preparation

We collected saliva samples from 24 unrelated Bouyei individuals in Bannong County in Guizhou Province. The participant’ parents and grandparents are indigenous people residing in the sampling palaces for at least three generations. All participants should have non-consanguineous marriages in the same ethnical group. The Medical Ethics Committees of West China Hospital of Sichuan University (2023-306) approved the study protocol. Besides, the procedure followed the recommendations of the Helsinki Declaration as revised in 2000. All the participants signed written informed consent before participating in the study.

### Relatedness analysis

We performed KING2 to test individual relationships by calculating kinship coefficients before merging data. All unrelated samples were kept for the following analyses.

### Data merging

We merged our genotyped Bouyei data with publicly available and previously published 38 TK-speaking populations from Southern China and SEA datasets, forming the primary TK datasets to explore the population structure and genetic relationship of whole TK-speaking populations(3, 32, 39, 47, 56). The TK datasets included 796 individuals from 38 populations, 522 from Southern China and 274 from SEA. Besides, to explore the genetic relationship between TK-speaking people and other modern and ancient eastern Eurasians, we merged our genome-wide data with the previously publicly modern and ancient people from Allen Ancient DNA Resource (AADR), which formed low-density merged HO dataset(26, 27), including 56814 SNPs; and middle-density merged 1240K dataset, including 146,802 SNPs. To explore the fine-scale genetic structure and obtain phased genomes to illuminate the biological adaptation mechanisms, we combined our data with previously published Illumina data(43–49), 54 worldwide populations included in the Human genetic diversity project (HGDP)(26), 20 populations from Taiwan Island, SEA Island, and Oceania included in the Oceanian genomic resources(27), which formed the high-density merged WGS dataset and included 460,678 SNPs.

### Principal Component Analysis

Using the merged Human Origin dataset, we carried out principal component analysis (PCA) via the smart PCA program of the EIGENSOFT v.6.1.4 package(56). All default parameters were used with the additional parameters of the lsq project: YES. Elsewise, we used PLINK to prune them with the parameters “-indep-pairwise 200 25 0.” (49). We performed three levels of PCA analyses, focusing on the relationship between different regional East Asian populations and the TK-speaking populations. PCA was first performed based on 207 modern and ancient East Asian populations to explore genetic similarities(57). Secondly, East Asian modern populations were extracted for further intra-regional PCA. The ancient samples from YRB, AMR_WLR, Guangxi, Fujian, Taiwan Island, and other language speakers were projected on the PCA analysis panel of the contemporary East Asians(58). We explored the genetic affinity of studied groups and other ethnic groups in 117 southern reference populations. Lastly, we performed a fine-scale analysis within 39 TK-speaking populations.

### Model-based ADMIXTURE analysis

The model-based maximum likelihood clustering algorithm ancestry-estimation method ADMIXTURE (59) was applied to explore the genetic composition using the merged Illumina, HGDP, and Oceanic datasets of 3514 individuals from 207 modern worldwide populations. We used ADMIXTURE mainly to identify Bouyei genetic structure and estimate individuals’ ancestry in the context of worldwide and regional people. Because the model in ADMIXTURE did not consider linkage disequilibrium, we used PLINK v.1.0711(49) to prune the original dataset with dense SNPs before analysing the ADMIXTURE. We ran ADMIXTURE 100 times with default parameters with the predefined ancestry sources or clusters, with the number of ancestral populations from K = 2 to K = 20 in bootstrap sequences with different random seeds (59). We selected an optimal K value based on assessing the lowest cross-validation error value and the highest log-likelihood using 10-fold cross-validation with different random seeds (60). Meanwhile, we performed ADMIXTURE containing 117 modern and ancient samples based on the low-density dataset to explore the genetic structure in an archaic context.

### Pairwise F_ST_ genetic distances

Pairwise F_ST_ (fixation index) accounted for genetic differences and evaluated the genetic affinity among different populations between these reference panels on the high-density dataset PLINK v1.90(5). Meanwhile, we calculated the pairwise F_ST_ genetic distance to measure the genetic relationship within Guizhou and other SEA TK-speaking populations based on the low-density dataset following Weir and Cockerham REFE 62 method.

### TreeMix analyses

We constructed the TreeMix-based phylogenetic tree among 14 populations to infer the genetic relationship and evaluate the gene flow events between Bouyei_Banong and other TK-speaking populations (61). Using the Illumina dataset array, a phylogenetic tree with migration events varying from 0 to 2 was reconstructed to study the genetic patterns of population split and admixture between our target populations and multiple ancestral populations.

### Runs of homozygosity (ROH)

We estimated the indicator of genomic homozygosity within 39 TK-speaking populations based on the low-density dataset using PLINK v1.90 (49). We set the ROH containing at least 50 SNPs, consecutive SNPs more than 100 kb apart, which were regarded as independent ROH. Lastly, we visualised the ROH distribution of each TK-speaking population statistically using R version 3.5.2 via the box plots. Further, we explore the length distribution patterns of ROH within TK-speaking people in Guizhou based on a high-density dataset.

### IBD Estimation and Effective Population Size

The IBD blocks were divided into three categories: −1, 1-5, and >5cM, which correspond very roughly to time intervals of early events (about1,500-2,500 years ago), interim events (about 500-1,500 years ago), and recent events (about 0-500 years ago), respectively. While considering the impact of impurities, we eliminated the smallest IBD segments, which reflected ancient events. Hence, we generated just two catalogues, 1-5, and >5cM using Refined-IBD software (16May19. ad5. jar) with a length parameter of 0.(62). Ne was used to estimate the effective population size among geographically diverse Bouyei people.

### Admixture-*f_3_*-Statistics and Outgroup-*f*_3_-Statistics

We used ADMIXTOOLS software(63) to compute *f*-statistic values and estimate standard errors by a block jackknife and default parameters. Firstly, we explored the potentially existing admixture signals within Bouyei and other modern/ancient populations in East Asia and performed the admixture-*f*_3_-statistics in the form *f*_3_(Source 1, Source 2; Targeted population) through the qp3pop to confirm that if the studied population was admixed. The target population with a negative *f*_3_ value and the value about |Z-score|>3 was regarded as a potential admixture signal. Since the *f_3_*-based estimates produce statistically significant values, we lowered the threshold, set the |Z-score|>2, and visualised the top 20 values used in R packages. Secondly, we selected modern East Asian and ancient populations as reference populations and performed the outgroup-*f*_3_ statistics in *f*_3_(Reference, studied populations; Mbuti) via the qp3Pop program of EIGENSOFT to explore the genetic affinity and drift between studied populations and other reference populations. Here, we conducted four group analyses, including 39 TK-speaking populations, 37 TK-speaking populations (not containing CentralThai and SouthernThai_TK), 42 ancient populations, and 38 Guizhou populations and visualised heatmap using R packages. A higher value and dark colour indicated a closer relationship.

### *F*_4_ Statistics

We conducted four population tests for targeted people based on the individual sample and merged populations (63). We calculated the *f*_4_-statistics to explore the signals and directions of admixture and the primary source of gene flow to Bouyei populations and other modern and ancient reference populations. Modern reference populations based on low-density datasets were used to test the genetic difference among the geographically different Bouyei people; ancient reference populations based on middle-density datasets were to be verified further the *f*_4_-statistics accuracy mentioned above. Therefore, we used qpDstat in ADMIXTOOLS to conduct the *f*_4_(Studied1, Studied2; reference populations, Mbuti) to explore the genetic heterozygosity and homogeneity among studied groups visualised the top 62 values used in R packages, here reference including ancient people based on middle-density dataset and modern people based on the low-density dataset. Then we performed the *f*_4_-statistics in the form of (Reference population1, Reference population2; BBN, Mbuti) to test genetic affinity. For ancient people, Reference population1 included eight ancient Northern China-YRB farmers of China_NEastAsia_Coastal_EN, Upper_YR_LN, himao_LN, YR_MN, YR_LN and YR_LBIA, and Reference population2 included ancient people from Guangxi, Fujian and SEA. For modern people, the reference populations included AA/AN/TK/HM-related southern populations and other northern populations. Elsewise, we merged four different geographical locations of Bouyei people as Bouyei populations to search for optimum ancestry sources on the population level in the form *f*_4_ (Reference populations, Meta-Bouyei; BBQC, Mbuti) and in the form *f*_4_ (BBQC, Meta-Bouyei; Reference populations, Mbuti). Reference populations included ancient people from different historical periods.

### Pairwise qpWave and qpAdm estimation

We used the qpAdm(64) with default parameters implemented in the ADMIXTOOLS package to estimate corresponding admixture proportions quantitatively. We used ancient northern East Asians as the northern surrogate of the ancestral source and used southern East Asians as the southern surrogate to model the formation of modern Bouyei people via qpAdm. We simulated the modern ancestral admixture model using northern Han, Tu, Hezhen, Xibo, Mongolian, Daur and Oroqen as the modern northern sources and used Li as the modern southern source. We used Yakut, Mbuti, Iran_GanjDareh_N, Villabruna, Ami, Mixe, Onge, Papuan and Ust_Ishim. Moreover, we performed the ancient ancestral admixture model using Miaozigou_MN, NEastAsia_Coastal_EN, NEastAsia_Inland_EN, Shimao_LN, Upper_YR_LN, WLR_LN, YR_LN, YR_MN, WLR_MN as the ancient northern sources and used China_SEastAsia_Island_LN, Taiwan_Hanben_IA, Yiyang, Layi as the ancient southern sources when Australian.DG, Russia_MA1_HG.SG as outgroups, respectively.

We also conducted pairwise qpWave analysis on the “merged-HO” dataset among TK/HM/AA/AN/TB, Sinitic, and Tungusic modern people to explore their genetic homogeneity within different pairwise populations and geographic scales. “+” indicated p_rank 0>0.01 and was statistically significant, “++” indicated p_rank 0>0.05. We re-performed pairwise qpWave analysis within form Northern and Southern ancient people to explore homogeneity and heterogeneity within the Bouyei population.

### ALDER Analysis

We used MALDER to estimate the generations, explore the admixture signatures, and fit the potential admixture time with the default parameters(65).

### CHROMOPAINTER and fastGLOBETROTTER

We used ChromoPainterv2 to paint the ancestral haplotype compositions and surrogates of four Bouyei populations. In this case, we ran fastGLOBETROTTER based on the default parameters to identify, describe, and date the admixture events(66).

### Painting Chromosomes and fineSTRUCTURE Analysis

We used SHAPEIT v2 (Segmented Haplotype Estimation & Imputation Tool) software with the default parameters (--burn 10 --prune 10 --main 30) to phase the genome-wide data of four Bouyei populations in Guizhou and other ten populations in geographically neighboring regions based on high-density merged WGS dataset and other 35 TK-speaking populations based on low-density merged HO dataset(67). Then, to dissect the fine-scale population stratifications, we conducted the ChromoPainter to compute the sharing haplotypes and obtain the coancestry matrix(66). Meanwhile, we used R packages implemented in the fineSTRUCTURE and matched the Admixture analysis to explore the phylogenetic relationship of Bouyei populations and fine-scale structure(66).

### Signatures of natural selection

We applied PBS to detect population and region-specific natural selection signals in Bouyei populations (68). Firstly, to explore the ancient natural selection signal, we used Northern Han (Shaaxi_Han) as an ingroup and European as an outgroup. Secondly, we also explored the regional natural selection using Bouyei_Guizhou as the target population and Zhuang/Shui/Mulam/Maonan/Gelao in Guizhou and Shaaxi_Han as the ingroup and outgroup, respectively. The top 0.1% values in PBS calculation were considered candidates, and the PBS calculation formula was: PBS_A_ = (T_AB_+T_AC_−T_BC_)/2, T= −log(1−F_ST_); where A as the target population, B and C are ingroups and outgroup population, respectively. In addition, based on the 10K_CPGDP, we calculated the allele frequency of selected alleles and mapped the frequency distribution in the globe.

### Functional Annotation of Natural Selection Signatures

We identified 141 and 132 candidates with variants of PBS values in the top 0.1% percentile in two pairs of population and regional-specific analyses, respectively. To search the candidates associated with different pathways in humans, we selected these candidates as the input gene set to perform functional enrichment by Metascape (https://metascape.org), which incorporated numerous functional categories and was beneficial for further analysis (69). In this analysis, we just used the following ontology sources: GO Biological Processes, Reactome Gene Sets, and WikiPathway to perform functional enrichment. The top 20 functional categories with −log10_(P-value)_ ≥2 was viewed as enriched pathways.

## Acknowledgments

This work was supported by grants from the National Natural Science Foundation of China (82202078), City-school science and technology strategic cooperation projects (22SXQT0351). We thank Prof. Mark Stoneking, Prof. Dang Liu at Max Planck Institute for Evolutionary Anthropology, and Prof. Wibhu Kutanan at Khon Kaen University for sharing genome-wide SNP data from Vietnam, Thailand, and Laos. We thank Prof. Etienne Patin and Prof. Lluis Quintana-Murci from the Human Evolutionary Genetics Unit of the Institute Pasteur for sharing the high-coverage genomes of 317 individuals from the Pacific region.

## Author Contributions

G.H. and J.Y. conceived and supervised the project. G.H., Y.L., and S.D. collected the samples. G.H., M.W., and Q.S. performed the extraction of the genomic DNA and coordinated the genome sequencing. S.D., Q.S., J.C., Z.W., Y.S., X.L., X.J., J.Y., C.W., C.L., M.W., G.H., and R.T. performed population genetic analysis. S.D., M.W., Y.L., and G.H. drafted the manuscript. G.H. and J.Y. revised the manuscript.

## Data Availability

The allele frequency data derived from human samples have been deposited in the National Omics Data Encyclopedia (NODE, http://www.biosino.org/node) and can be accessed with accession number (OEPXXXXXX, available after publication). The access and use of the data shall comply with the regulations of the People’s Republic of China on the administration of human genetic resources. Requests for access to data can be directed to Guanglin He.

## Ethics declarations

### Ethics approval and consent to participate

The Medical Ethics Committees of West China Hospital of Sichuan University approved this study. This study was conducted by the principles of the Helsinki Declaration.

### Consent for publication

Not applicable.

## Reference

1. Yang MA, Fan X, Sun B, Chen C, Lang J, Ko YC, et al. Ancient DNA indicates human population shifts and admixture in northern and southern China. Science. 2020;369(6501):282–8.

2. Mao X, Zhang H, Qiao S, Liu Y, Chang F, Xie P, et al. The deep population history of northern East Asia from the Late Pleistocene to the Holocene. Cell. 2021;184(12):3256–66 e13.

3. Wang CC, Yeh HY, Popov AN, Zhang HQ, Matsumura H, Sirak K, et al. Genomic insights into the formation of human populations in East Asia. Nature. 2021;591(7850):413–9.

4. Wang H, Yang MA, Wangdue S, Lu H, Chen H, Li L, et al. Human genetic history on the Tibetan Plateau in the past 5100 years. Science advances. 2023;9(11):eadd5582.

5. Lipson M, Cheronet O, Mallick S, Rohland N, Oxenham M, Pietrusewsky M, et al. Ancient genomes document multiple waves of migration in Southeast Asian prehistory. Science. 2018;361(6397):92–5.

6. McColl H, Racimo F, Vinner L, Demeter F, Gakuhari T, Moreno-Mayar JV, et al. The prehistoric peopling of Southeast Asia. Science. 2018;361(6397):88–92.

7. Ning C, Li T, Wang K, Zhang F, Li T, Wu X, et al. Ancient genomes from northern China suggest links between subsistence changes and human migration. Nat Commun. 2020;11(1):2700.

8. Diamond J, Bellwood P. Farmers and their languages: the first expansions. Science. 2003;300(5619):597–603.

9. WANG C-C. The Genomic Formation of Human Populations in East Asia. 2020.

10. jin L. Genetic, Linguistic and Archaeological Perspectives on Human Diversity in Southeast Asia. 2001.

11. Gyaneshwer Chaubey1 MM, Ying Choi2, Reedik Mägi3,4, Irene Gallego, Romero5 PS, Mannis van Oven2, Doron M. Behar1,7, Siiri Rootsi1, Georgi, Hudjashov1 CBM, Monika Karmin1, Mari Nelis8, Jüri Parik1, Alla, Goverdhana Reddy9 EM, George van Driem10, Yali Xue11, Chris Tyler-Smith11,, Kumarasamy Thangaraj9 LS, Maido Remm4, Martin B. Richards6, Marta Mirazon, Lahr5 MK, Richard Villems1, and Toomas Kivisild. Population Genetic Structure in Indian Austroasiatic Speakers: The Role of Landscape Barriers and Sex-Specific Admixture.

12. He G, Ren Z, Guo J, Zhang F, Zou X, Zhang H, et al. Population genetics, diversity and forensic characteristics of Tai-Kadai-speaking Bouyei revealed by insertion/deletions markers. Molecular genetics and genomics: MGG. 2019;294(5):1343–57.

13. He G, Wang Z, Guo J, Wang M, Zou X, Tang R, et al. Inferring the population history of Tai-Kadai-speaking people and southernmost Han Chinese on Hainan Island by genome-wide array genotyping. Eur J Hum Genet. 2020;28(8):1111–23.

14. Xiaoming-Zhang. A Matrilineal Genetic Perspective of Hanging Coffin Custom in Southern China and Northern Thailand.

15. Wang T, Wang W, Xie G, Li Z, Fan X, Yang Q, et al. Human population history at the crossroads of East and Southeast Asia since 11,000 years ago. Cell. 2021;184(14):3829–41 e21.

16. Chen H, Lin R, Lu Y, Zhang R, Gao Y, He Y, et al. Tracing Bai-Yue Ancestry in Aboriginal Li People on Hainan Island. Mol Biol Evol. 2022;39(10):msac210.

17. Sun J, Li YX, Ma PC, Yan S, Cheng HZ, Fan ZQ, et al. Shared paternal ancestry of Han, Tai-Kadai-speaking, and Austronesian-speaking populations as revealed by the high resolution phylogeny of O1a-M119 and distribution of its sub-lineages within China. Am J Phys Anthropol. 2021;174(4):686–700.

18. Kutanan W, Kampuansai J, Fuselli S, Nakbunlung S, Seielstad M, Bertorelle G, et al. Genetic structure of the Mon-Khmer speaking groups and their affinity to the neighbouring Tai populations in Northern Thailand. BMC genetics. 2011;12:56.

19. Kutanan W, Kampuansai J, Srikummool M, Kangwanpong D, Ghirotto S, Brunelli A, et al. Complete mitochondrial genomes of Thai and Lao populations indicate an ancient origin of Austroasiatic groups and demic diffusion in the spread of Tai-Kadai languages. Hum Genet. 2017;136(1):85–98.

20. Kutanan W, Ghirotto S, Bertorelle G, Srithawong S, Srithongdaeng K, Pontham N, et al. Geography has more influence than language on maternal genetic structure of various northeastern Thai ethnicities. J Hum Genet. 2014;59(9):512–20.

21. Srithawong S, Srikummool M, Pittayaporn P, Ghirotto S, Chantawannakul P, Sun J, et al. Genetic and linguistic correlation of the Kra-Dai-speaking groups in Thailand. J Hum Genet. 2015;60(7):371–80.

22. Deng QY, Wang CC, Wang XQ, Wang LX, Wang ZY, Wu WJ, et al. Genetic affinity between the Kam-Sui speaking Chadong and Mulam people. Journal of Systematics and Evolution. 2013;51(3):263–70.

23. Wang MG, He GL, Zou X, Chen PY, Wang Z, Tang RK, et al. Reconstructing the genetic admixture history of Tai-Kadai and Sinitic people: Insights from genome-wide SNP data from South China. Journal of Systematics and Evolution. 2022;61(1):157–78.

24. He G, Wang Z, Zou X, Wang M, Liu J, Wang S, et al. Tai-Kadai-speaking Gelao population: Forensic features, genetic diversity and population structure. Forensic Sci Int Genet. 2019;40:e231–e9.

25. Ren Z, Guo J, He G, Zhang H, Zou X, Zhang H, et al. Forensic genetic polymorphisms and population structure of the Guizhou Bouyei people based on 19 X-STR loci. Ann Hum Biol. 2019;46(7-8):574–80.

26. Bergstrom A, McCarthy SA, Hui R, Almarri MA, Ayub Q, Danecek P, et al. Insights into human genetic variation and population history from 929 diverse genomes. Science. 2020;367(6484).

27. Choin J, Mendoza-Revilla J, Arauna LR, Cuadros-Espinoza S, Cassar O, Larena M, et al. Genomic insights into population history and biological adaptation in Oceania. Nature. 2021;592(7855):583–9.

28. Fumagalli M, Moltke I, Grarup N, Racimo F, Bjerregaard P, Jorgensen ME, et al. Greenlandic Inuit show genetic signatures of diet and climate adaptation. Science. 2015;349(6254):1343–7.

29. Qiaomei-Fu. Human population history at the crossroads of East and Southeast Asia since 11,000 years ago.

30. Kosoy R, Ransom M, Chen H, Marconi M, Macciardi F, Glorioso N, et al. Evidence for malaria selection of a CR1 haplotype in Sardinia. Genes and immunity. 2011;12(7):582–8.

31. Pascale Gaudet MSL, Suzanna E. Lewis and Paul D. Thomas. Phylogenetic-based propagation of functional annotations within the Gene Ontology consortium.

32. Kutanan W, Liu D, Kampuansai J, Srikummool M, Srithawong S, Shoocongdej R, et al. Reconstructing the Human Genetic History of Mainland Southeast Asia: Insights from Genome-Wide Data from Thailand and Laos. Mol Biol Evol. 2021;38(8):3459–77.

33. Gaspard Kerner A-LN, Quentin Philippot, …, Etienne Patin, Guillaume Laval, Lluis Quintana-Murci. Genetic adaptation to pathogens and increased risk of inflammatory disorders in post-Neolithic Europe.

34. Ian A. Cockburn* MJM, Angela O’Donnell†, Stephen J. Allen†‡, Joann M. Moulds§, Moses Baisor¶,, Moses Bockarie¶ JCR, aJAR. A human complement receptor 1 polymorphism that reduces Plasmodium falciparum rosetting confers protection against severe malaria.

35. Kosoy1 R, MR, HC, MM, FM, 5, N Glorioso6, PG, et al. Evidence for malaria selection of a CR1 haplotype in Sardinia.

36. Vandana Thathy1 JMM, 4, Bernard Guyah1, Walter Otieno1 and José A Stoute. Complement receptor 1 polymorphisms associated with resistance to severe malaria in Kenya.

37. Mengge W, Guanglin H, Yongdong S, Shouyu W, Xing Z, Jing L, et al. Massively parallel sequencing of mitogenome sequences reveals the forensic features and maternal diversity of tai-kadai-speaking hlai islanders. Forensic Sci Int Genet. 2020;47:102303.

38. Zhao ZJ. New Archaeobotanic Data for the Study of the Origins of Agriculture in China. Current Anthropology. 2011;52(S4):S295–S306.

39. Liu D, Duong NT, Ton ND, Van Phong N, Pakendorf B, Van Hai N, et al. Extensive Ethnolinguistic Diversity in Vietnam Reflects Multiple Sources of Genetic Diversity. Mol Biol Evol. 2020;37(9):2503–19.

40. Kutanan W, Kampuansai J, Srikummool M, Brunelli A, Ghirotto S, Arias L, et al. Contrasting Paternal and Maternal Genetic Histories of Thai and Lao Populations. Mol Biol Evol. 2019;36(7):1490–506.

41. Peng MS, He JD, Liu HX, Zhang YP. Tracing the legacy of the early Hainan Islanders--a perspective from mitochondrial DNA. BMC Evol Biol. 2011;11(1):46.

42. Li D-N, Wang C-C, Lu Y, Qin Z-D, Yang K, Lin X-J, et al. Three phases for the early peopling of Hainan Island viewed from mitochondrial DNA. Journal of Systematics and Evolution. 2013;51(6):671–80.

43. Wang Y, Zou X, Wang M, Yuan D, Yang L, Zeng Y, et al. The genomic history of southwestern Chinese populations demonstrated massive population migration and admixture among proto-Hmong-Mien speakers and incoming migrants. Molecular genetics and genomics: MGG. 2022;297(1):241–62.

44. He GL, Li YX, Zou X, Yeh HY, Tang RK, Wang PX, et al. Northern gene flow into southeastern East Asians inferred from genome-wide array genotyping. Journal of Systematics and Evolution. 2022;61(1):179–97.

45. He G, Zhang Y, Wei LH, Wang M, Yang X, Guo J, et al. The genomic formation of Tanka people, an isolated “gypsies in water” in the coastal region of Southeast China. American Journal of Biological Anthropology. 2022;178(1):154–70.

46. He G, Fan ZQ, Zou X, Deng X, Yeh HY, Wang Z, et al. Demographic model and biological adaptation inferred from the genome-wide single nucleotide polymorphism data reveal tripartite origins of southernmost Chinese Huis. American Journal of Biological Anthropology. 2022;180(3):488–505.

47. Chen J, He G, Ren Z, Wang Q, Liu Y, Zhang H, et al. Fine-Scale Population Admixture Landscape of Tai-Kadai-Speaking Maonan in Southwest China Inferred From Genome-Wide SNP Data. Front Genet. 2022;13:815285.

48. Zhang X, He G, Li W, Wang Y, Li X, Chen Y, et al. Genomic Insight Into the Population Admixture History of Tungusic-Speaking Manchu People in Northeast China. Front Genet. 2021;12(1761):754492.

49. Chang CC, Chow CC, Tellier LC, Vattikuti S, Purcell SM, Lee JJ. Second-generation PLINK: rising to the challenge of larger and richer datasets. Gigascience. 2015;4:7.

50. Wu S, Tan J, Yang Y, Peng Q, Zhang M, Li J, et al. Genome-wide scans reveal variants at EDAR predominantly affecting hair straightness in Han Chinese and Uyghur populations. Hum Genet. 2016;135(11):1279–86.

51. Kamberov YG, Wang S, Tan J, Gerbault P, Wark A, Tan L, et al. Modeling recent human evolution in mice by expression of a selected EDAR variant. Cell. 2013;152(4):691–702.

52. Kimura R, Yamaguchi T, Takeda M, Kondo O, Toma T, Haneji K, et al. A common variation in EDAR is a genetic determinant of shovel-shaped incisors. Am J Hum Genet. 2009;85(4):528–35.

53. Peng Q, Li J, Tan J, Yang Y, Zhang M, Wu S, et al. EDARV370A associated facial characteristics in Uyghur population revealing further pleiotropic effects. Hum Genet. 2016;135(1):99–108.

54. Zhihua Li. YP, Edward M. Marcotte. A Bacteriophage Tailspike Domain Promotes SelfCleavage of a Human Membrane-Bound Transcription Factor, the Myelin Regulatory Factor MYRF.

55. Jie Zhang JH, Xiao-Hong Zeng, Shi-Jun Ge, Yu Huang, Jie Su, Xue-Mei Ding, Ji-Qing Yang, Yong-Jiu Cao, Hong Chen, Ying-Hong Zhang,Bao-Sheng Zhu. Genetic Heterogeneity of the β-Globin Gene in Various Geographic Populations of Yunnan in Southwestern China.

56. Patterson N, Price AL, Reich D. Population structure and eigenanalysis. PLoS Genet. 2006;2(12):e190.

57. Choin J, Mendoza-Revilla J, Arauna LR, Cuadros-Espinoza S, Cassar O, Larena M, et al. Genomic insights into population history and biological adaptation in Oceania. Nature. 2021;592(7855):583–9.

58. McColl H, Racimo F, Vinner L, Demeter F, Gakuhari T, Moreno-Mayar JV, et al. The prehistoric peopling of Southeast Asia. Science. 2018;361(Jul.6 TN.6397):88–92.

59. Alexander DH, Novembre J, Lange K. Fast model-based estimation of ancestry in unrelated individuals. Genome Res. 2009;19(9):1655–64.

60. Feng Q, Lu D, Xu S. AncestryPainter: A Graphic Program for Displaying Ancestry Composition of Populations and Individuals. Genomics, Proteomics & Bioinformatics. 2018;16(5):382–5.

61. Pickrell JK, Pritchard JK. Inference of population splits and mixtures from genome-wide allele frequency data. PLoS Genet. 2012;8(11):e1002967.

62. Browning BL, Browning SR. Improving the accuracy and efficiency of identity-by-descent detection in population data. Genetics. 2013;194(2):459–71.

63. Patterson N, Moorjani P, Luo Y, Mallick S, Rohland N, Zhan Y, et al. Ancient admixture in human history. Genetics. 2012;192(3):1065–93.

64. Harney E, Patterson N, Reich D, Wakeley J. Assessing the performance of qpAdm: a statistical tool for studying population admixture. Genetics. 2021;217(4).

65. Loh PR, Lipson M, Patterson N, Moorjani P, Pickrell JK, Reich D, et al. Inferring admixture histories of human populations using linkage disequilibrium. Genetics. 2013;193(4):1233–54.

66. Lawson DJ, Hellenthal G, Myers S, Falush D. Inference of population structure using dense haplotype data. PLoS Genet. 2012;8(1):e1002453.

67. O.D. Jma. Improved whole-chromosome phasing for disease and population genetic studies.

68. Weir BS, Cockerham CC. Estimating F-Statistics for the Analysis of Population Structure. Evolution. 1984;38(6):1358–70.

69. Zhou Y, Zhou B, Pache L, Chang M, Khodabakhshi AH, Tanaseichuk O, et al. Metascape provides a biologist-oriented resource for the analysis of systems-level datasets. Nat Commun. 2019;10(1):1523.

