## Supplemental files for "Malaria resistance-related biological adaptation and complex evolutionary footprints of Tai-Kadai people inferred from 796 genomes"

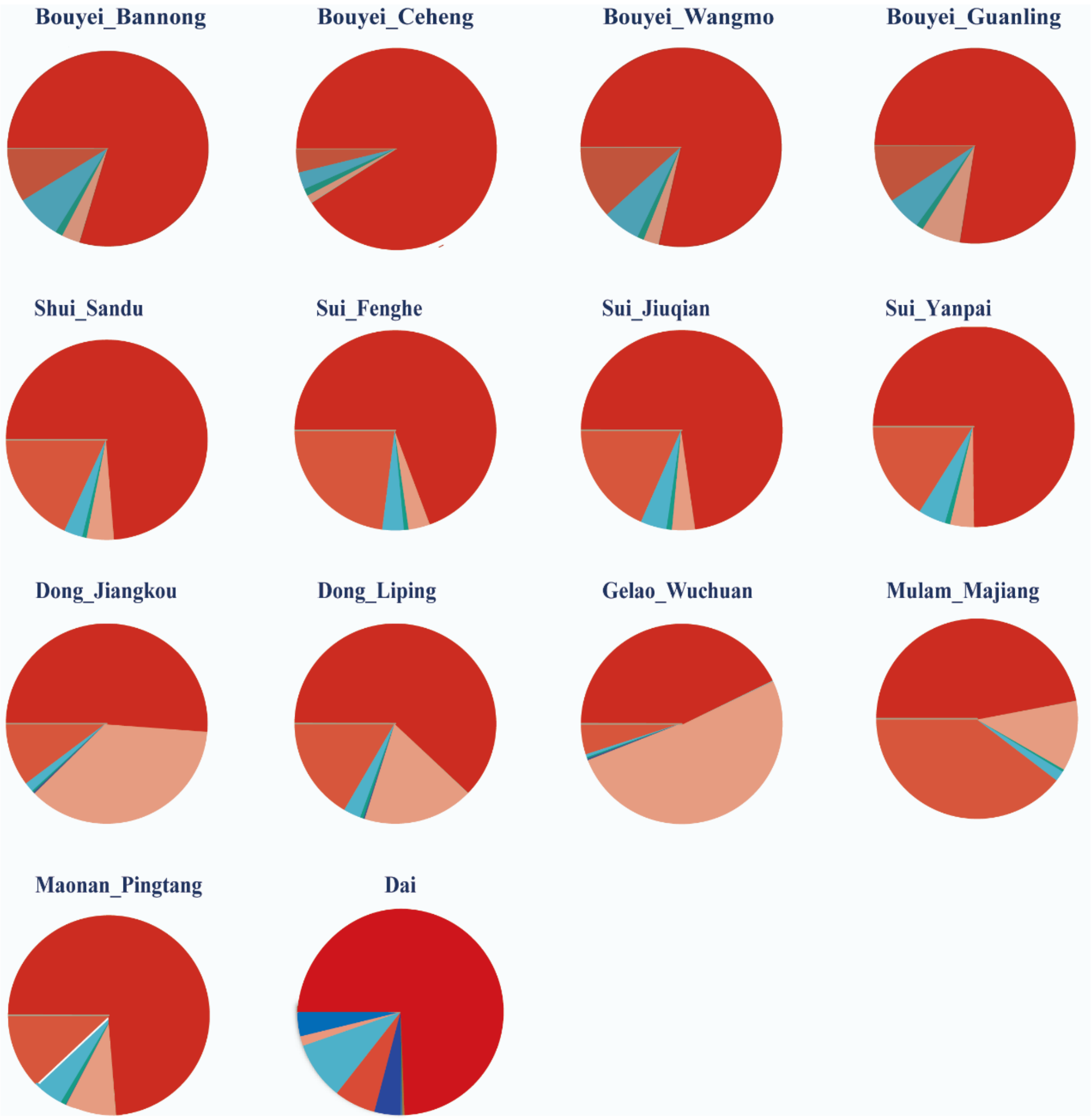


**Figure S1. The ancestry composition of fourteen TK populations in Guizhou.** Pies showed the proportions of each genetic component for 14 TK populations in Guizhou with 11 predefined ancestral sources.

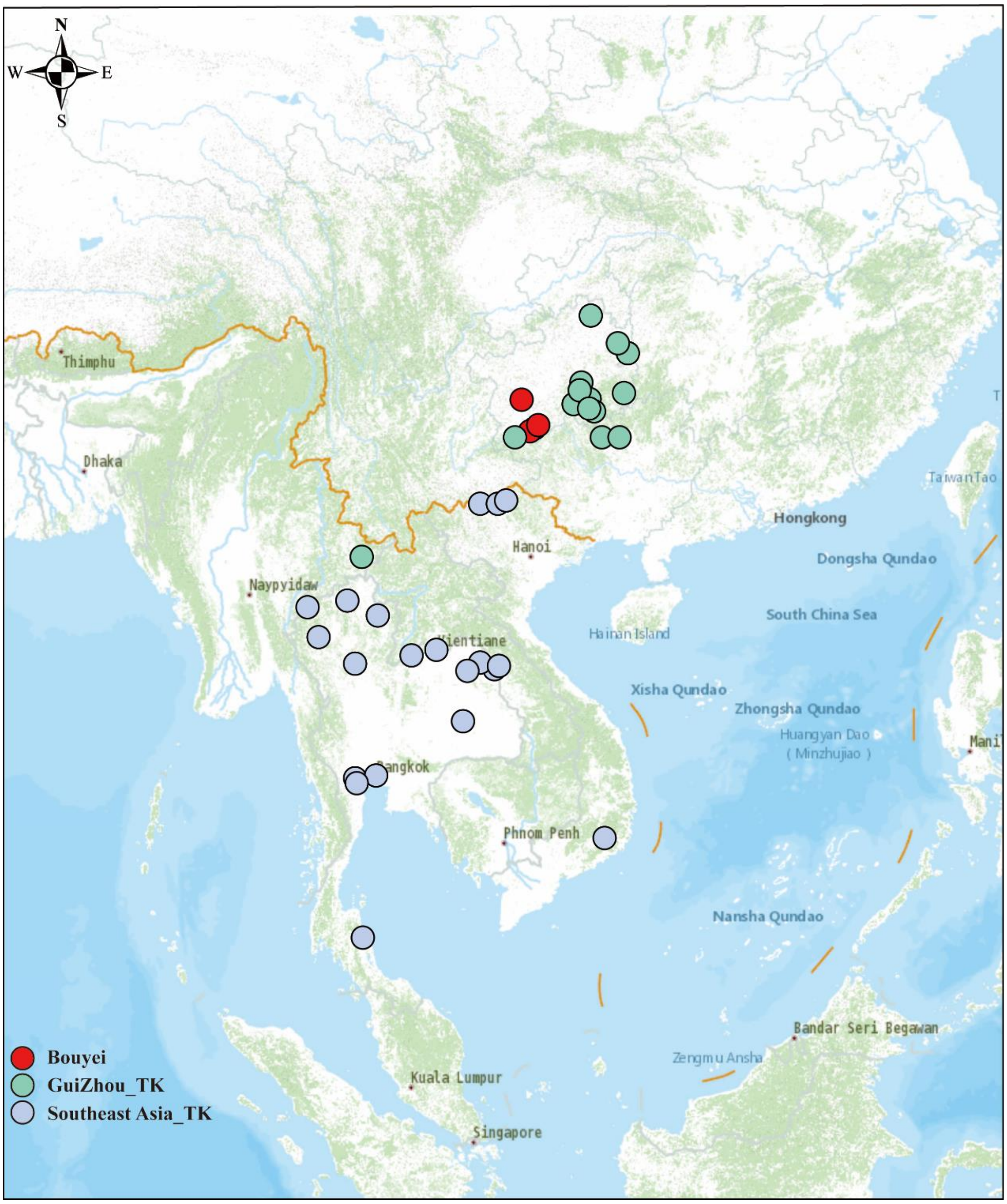


**Figure S2. Sample location of the 39 TK populations.**

The geographic locations of 39 TK populations included Southern China and MSEA in the low-density SNP dataset. Red circles represented four Bouyei populations; green circles represented 14 TK populations in Guizhou; blue circles represented 21 other populations from SEA.

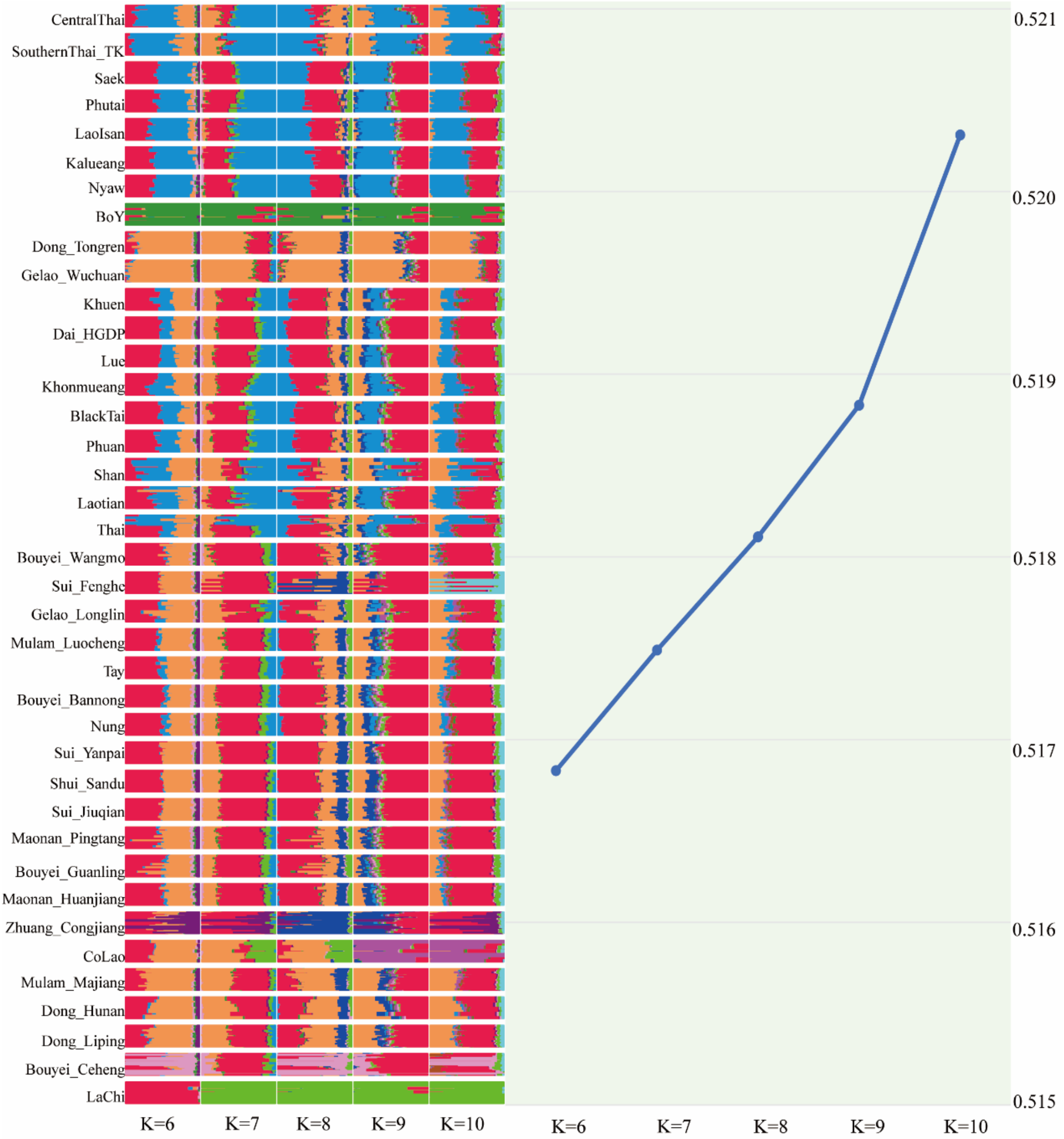


**Figure S3. Model-based ADMIXTURE analysis of the 39 TK populations with the 6–10 ancestral sources.**

Each increase in the value of K between 6 and 10 resulted in a single population being distinguished. On the right was a line graph of the cross-error values.

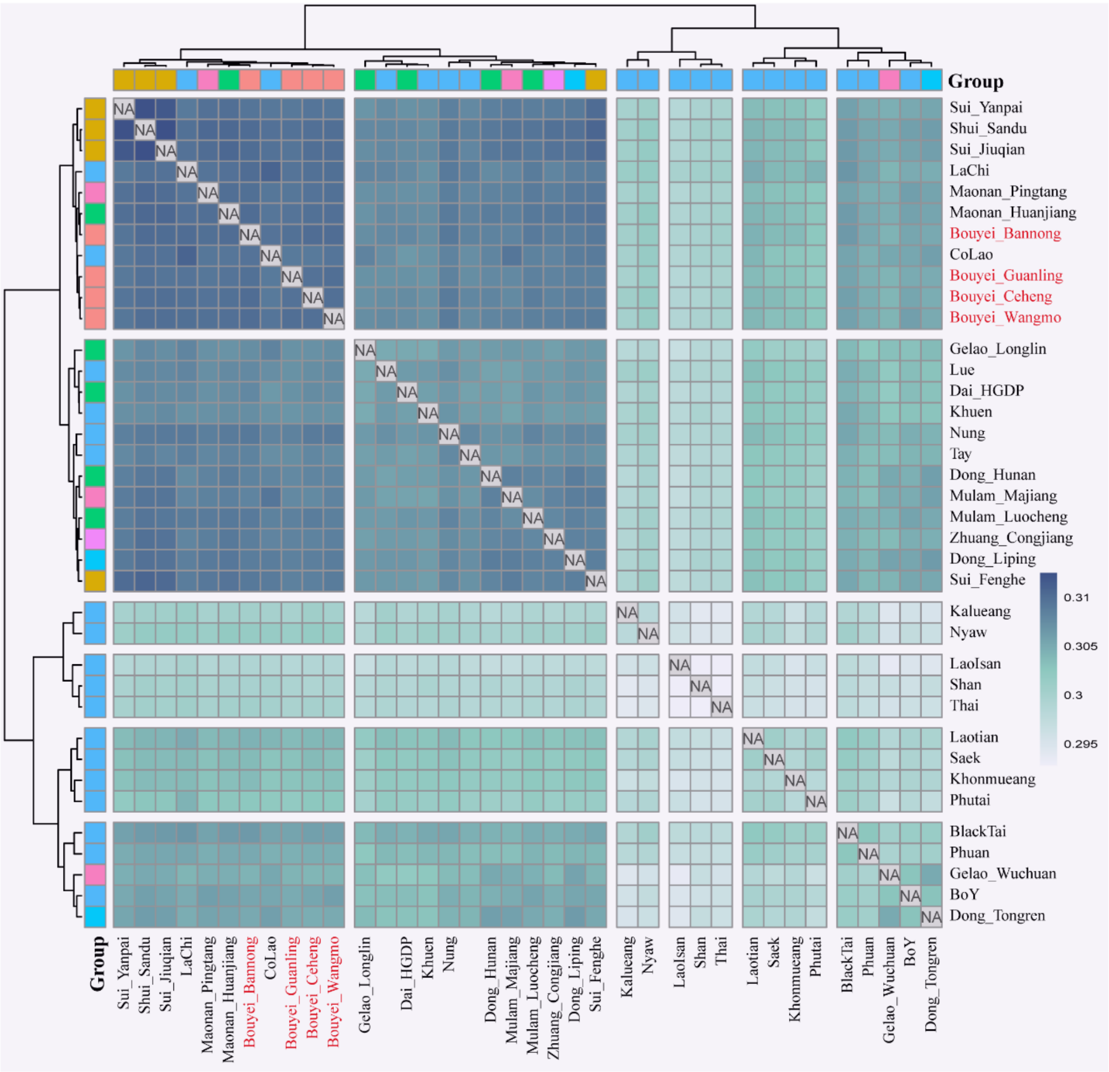


**Figure S4. Genetic affinities of the Bouyei populations based on the Outgroup-$f_3$ and shared genetic drift.**

Heatmap of the shared genetic drift inferred from Outgroup $f_3$-statistics in the form $f_3$(Studied Bouyei, TK; Mbuti), TK including 37 populations in Southern China and SEA except for CentralThai and SouthernThai_TK in the merged HO dataset, the genetic affinity between populations gradually increased from shallow to deep.

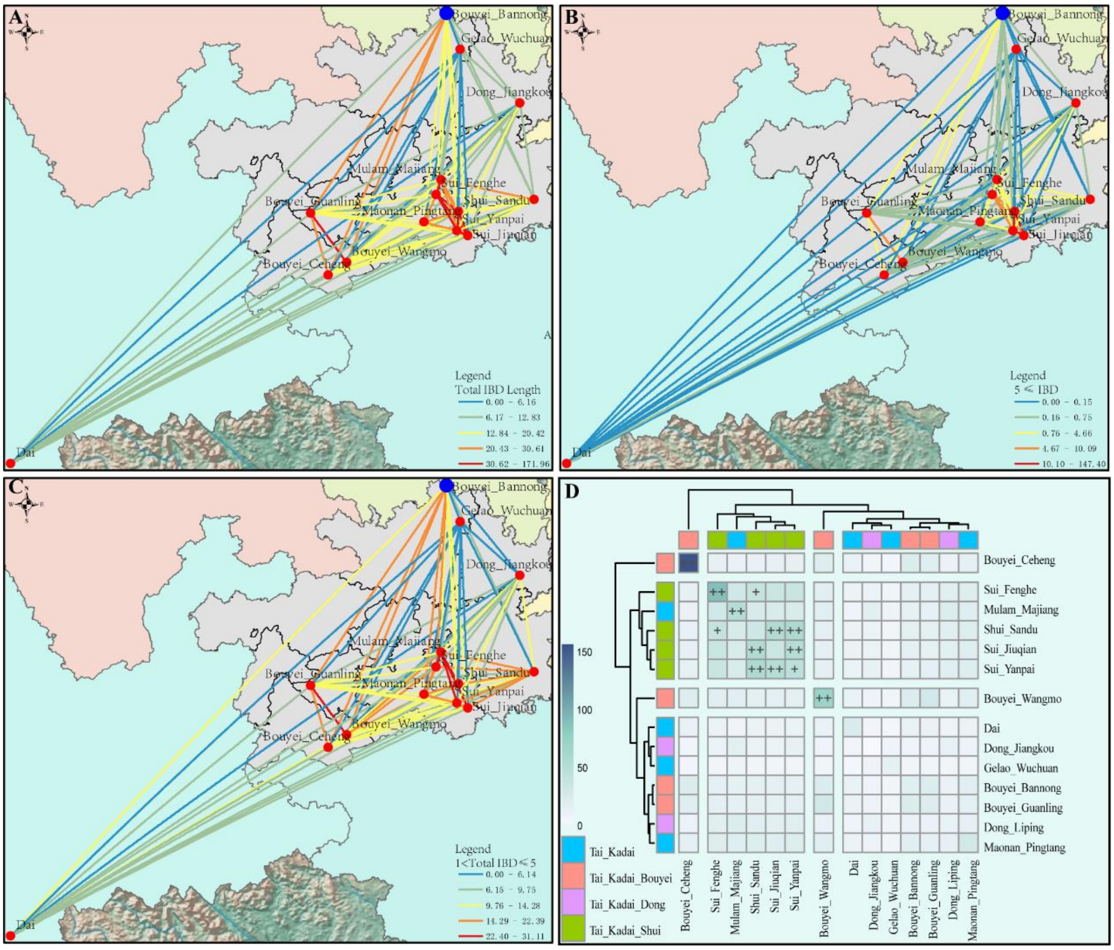


**Figure S5. Pairwise identity-by-descent (IBD) networks within the 14 populations in Guizhou.**

Shared identity by descent (IBD) fragments in different length ranges of Bouyei and other TK populations. Different colours represented the different total number of shared IBD fragments between populations. (**A-C**) (**A**)The total IBD length in 14 TK populations in Guizhou; (**B**) The pairwise IBD sharing average length between 14 populations more than 5; (**C**) The pairwise IBD sharing average length between 14 populations more than 1 but less than 5; (**D**) The pairwise IBD sharing average length between 14 populations all.

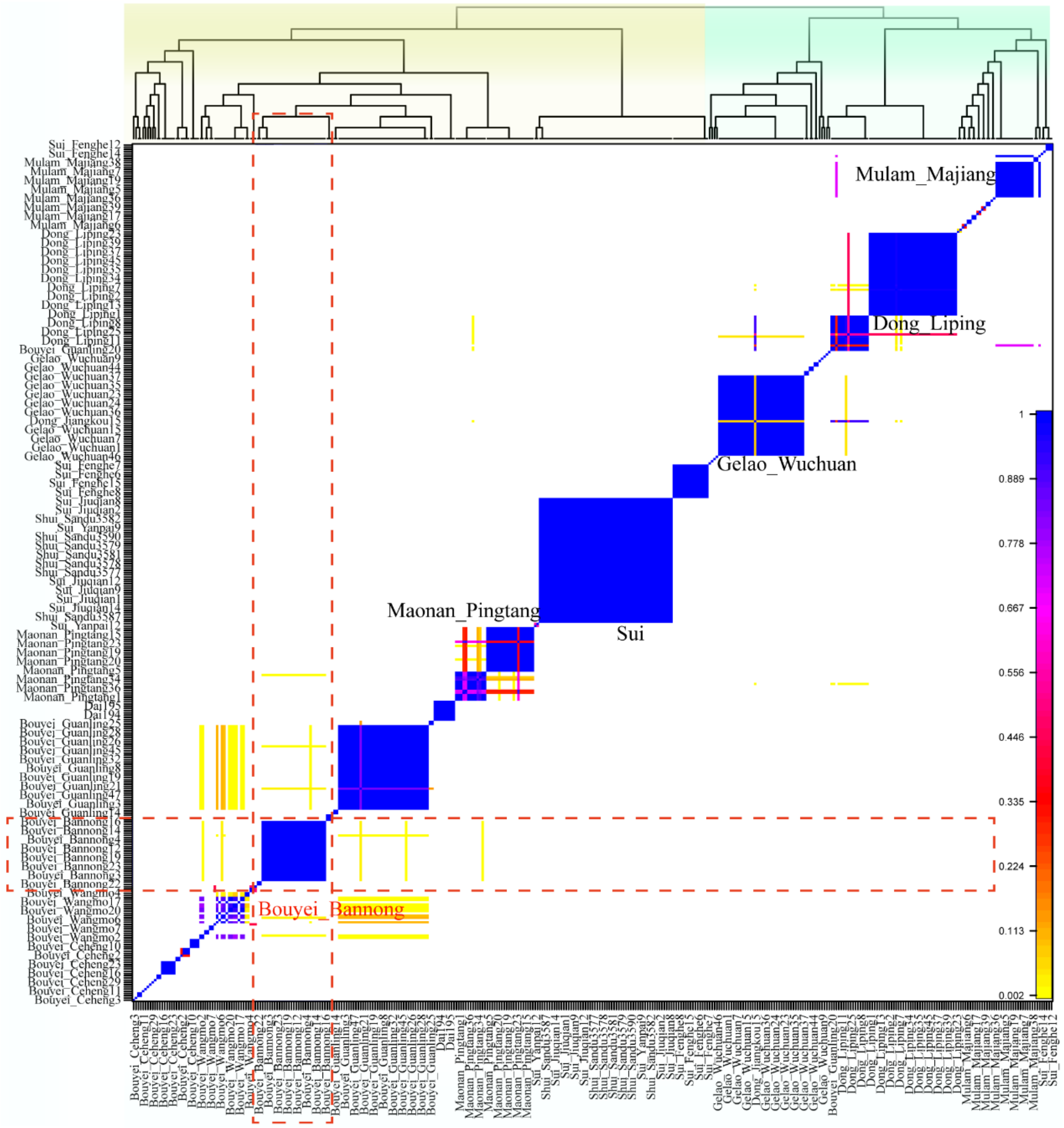


**Figure S6: Clustering patterns of TK individuals in Guizhou based on the pairwise coincidence matrix.**

Pairwise coincidence matrix at individual level inferred by fineSTRUCTURE with extremely high (dark blue) and low (yellow) values. In general, it can be divided into two clusters from FullDendrogram, and target groups (Bouyei_Bannong) were marked in red.

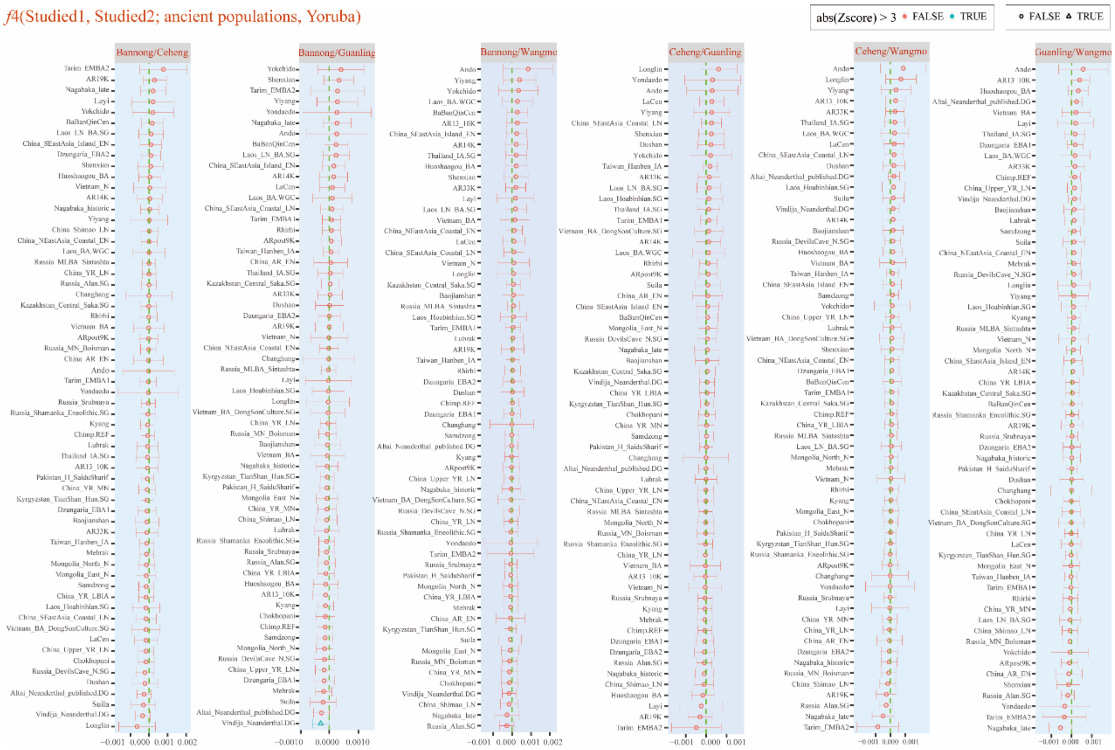

**Figure S7: $F_4$-statistic test for homogeneity and heterogeneity in four Bouyei groups.**

To test the genetic homogeneity and heterogeneity between targeted groups, $f_4$-statistics in the form of $f_4$(Studied1, Studied2; ancient populations, Mbuti). The |Z |values of $f_4$ under 3 standard errors indicated four Bouyei groups are genetic homogeneity.

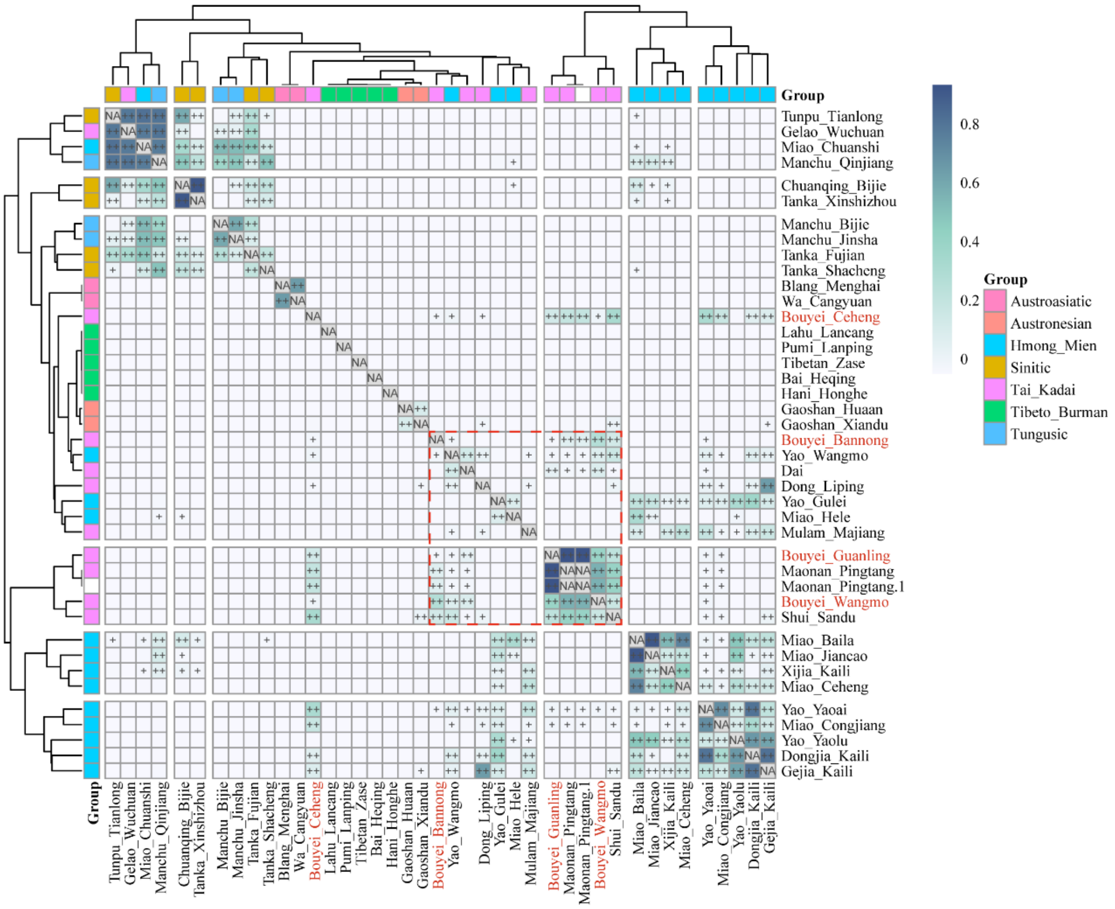


**Figure S8: Pairwise qpWave test for homogeneity and heterogeneity between Bouyei and other East Asian populations.**

Conducted pairwise qpWave analysis among 2 AA-, 2 AN-, 12 HM-, 5 Sinitic-, 5 TB-, 3 Tungusic- and 12 TK-speaking populations, “+” indicated p_rank0>0.01 and was statistically significant, ++ indicated p_rank0>0.05. The red rectangle showed the strong genetic homogeneity that existed within geographically and linguistically close TK populations in Guizhou.

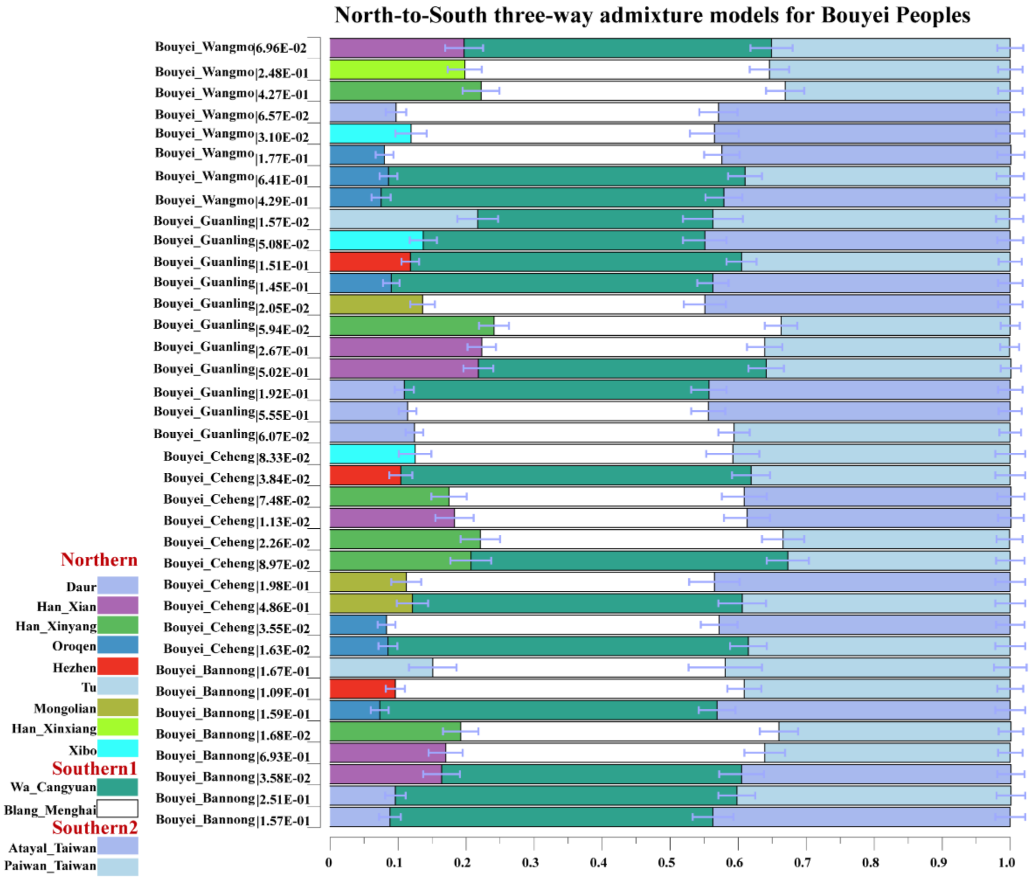


**Figure S9: Admixture proportion of ancestry groups from modern inferred using qpAdm.**

Three-way admixture models also showed both modern Northern and Southern populations contributed to the formation of four Bouyei people. The error bar indicated the stand errors of predicted proportions of ancestors obtained from qpAdm.

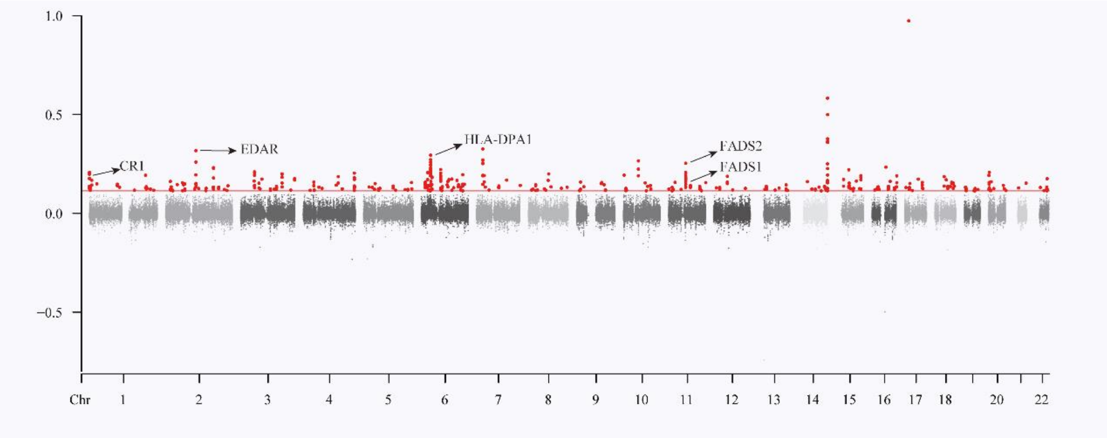


**Figure S10: Positive natural selection signals.**

Manhattan plot showing the PBS values in genome-wide scanned for Bouyei population in Guizhou, using the Shaanxi_Han and European as ingroup and outgroup reference populations. The 99.9th percentiles of the PBS distribution were shown as red lines. PBS values over the 99.9th percentile were marked in red, and PBS values under the 99.9th percentile were coloured as dark dots. Elsewise, some of the genes were labelled with its name.
